# Dual CRISPRi-Seq for genome-wide genetic interaction studies identifies key genes involved in the pneumococcal cell cycle

**DOI:** 10.1101/2024.08.14.607739

**Authors:** Julien Dénéréaz, Elise Eray, Bimal Jana, Vincent de Bakker, Horia Todor, Tim van Opijnen, Xue Liu, Jan-Willem Veening

## Abstract

A major goal in biology is to uncover the relationship between genotype and phenotype. However, identifying gene function is often hampered by genetic redundancy. For example, under standard laboratory conditions, three-quarters of the genes in the human pathogen *Streptococcus pneumoniae* are non-essential. A powerful approach to unravel genetic redundancy is by identifying gene-gene interactions. To uncover genetic interactions (GIs) in *S. pneumoniae* on a genome-wide scale, a generally applicable dual CRISPRi-Seq method and associated analysis pipeline was developed. Specifically, we created a library of 869 dual sgRNAs targeting high-confidence operons that encode essential and non-essential genes, covering over 70% of the genetic elements in the pneumococcal genome. Testing these 378,015 unique combinations, 4,026 significant GIs were identified, including 1,935 negative and 2,091 positive interactions. Besides known GIs, we found and confirmed previously unknown interactions involving genes responsible for fundamental cellular processes such as cell division, cell shape maintenance, and chromosome segregation. The presented methods and bioinformatic approaches can serve as a roadmap for genome-wide gene interaction studies in other organisms. Lastly, all interactions are available for exploration via the Pneumococcal Genetic Interaction Network (PneumoGIN) at https://veeninglab.shinyapps.io/PneumoGIN, which can serve as a starting point for new biological discoveries and translational research.

## Introduction

Understanding the relationship between genotype and phenotype is a central objective in biology. However, identifying gene function is challenging because not all genes exhibit a phenotype when disrupted. Although genome-scale genomics methods like transposon insertion sequencing (Tn-Seq) and CRISPRi-seq have made significant progress in this area^1,2^, many conserved genes remain of unknown function due to the absence of observable phenotypes upon disruption. This may be due to the inadequate representation of ecological conditions when testing mutant libraries, and/or due to partial redundancy between genes and pathways. Thus, determining the functions of these seemingly dispensable genes remains a major challenge. One avenue to address this problem is by genetic interaction (GI) studies. For instance, high-throughput synthetic genetic arrays (SGA) in *Saccharomyces cerevisiae* have enabled GI screens that have revealed the identification of protein complexes and many redundant genetic pathways^3–6^. A similar approach has been developed for *Escherichia coli* in which SGAs were used to map GIs by the pairwise conjugation of gene deletions^7^. In both cases, GIs are assessed by measuring colony sizes of the double knockouts, in which a complete lack of colony formation indicates a strong negative GI (synthetic lethality). Although these array-based approaches demonstrate their potency and underscore the significance of creating interaction networks, their applicability remains limited to a specific set of model organisms and environmental conditions that can be tested. Moreover, out of necessity, these tools mainly focus on GIs between non-essential genes, although essential genes can be queried by using temperature-sensitive or hypomorphic alleles^3,7,8^.

In the human pathogen *Streptococcus pneumoniae* (the pneumococcus), extensive screenings using Tn-Seq have shown that roughly three-quarters of its genes are non-essential under any given condition^9–11^. This suggests that many biological processes in this bacterium involve redundant genes, potentially acting as a fail-safe mechanism. Interrogating genetic interactions can highlight such mechanisms and help elucidate the underlying gene functions. Tn-Seq has mostly been used in different genetic backgrounds to assess genetic interactions. By comparing the fitness of each Tn-disrupted gene in a wild-type and query gene-knockout background, new functions and relationships among genes have been identified^10–13^. However, Tn-Seq requires a large library to fully cover the genome, which would scale substantially if genetic interactions were scored genome-wide^14^. More importantly, the major drawback of methods such as Tn-Seq and SGA methods is their inability to study essential genes, as knockouts are non-viable. Recently, a combination of CRISPR interference and Tn-Seq (CRISPRi-TnSeq) was used to investigate GIs of a dozen selected essential genes^15^. However, the resulting genetic interaction network was limited to those selected essential genes targeted by CRISPRi and the non-essential genes targeted by Tn-Seq.

To overcome these limitations, we developed dual CRISPR interference (dual CRISPRi-Seq) to investigate GIs on a genome-wide scale. CRISPRi relies on a target-specific single guide RNA (sgRNA) and an inducible catalytically dead Cas9 (dCas9) which is guided to the targeted gene, thus acting as a roadblock for RNA polymerase repressing transcription^16,17^. CRISPRi operates at the operon level due to its polar effects and can target both non-essential and essential genes. Single CRISPRi has been instrumental in the identification of genes involved in pneumococcal cell division, competence, virulence, and cell wall synthesis^9,18,19^. Dual CRISPRi has hitherto only been performed at a large scale in human cell lines by knocking down genes using two different sgRNAs^4,5,20,21^. The potential of this approach was demonstrated by highlighting interdependency between core pathways like cholesterol biosynthesis and DNA repair, and the identification of potential targets for anti-cancer combination therapy^4^. Furthermore, by utilizing the native *Streptococcus pyogenes* CRISPR array, multiplex CRISPRi was used to repress the transcription of multiple genes in *Legionella pneumophila* and *Escherichia coli* simultaneously^22–24^. However, a genome-wide method to assess genetic interactions by CRISPRi has not yet been realized for bacteria.

We previously engineered a genome-wide pooled CRISPRi library in *S. pneumoniae* D39V, which, combined with next-generation sequencing (NGS), allows for genome-wide fitness quantifications of all targeted genes under a chosen condition^1,25^. Building on CRISPRi-Seq, a generally applicable approach to study GIs on a genome-wide level called dual CRISPRi-seq is presented here. To this end, a library composed of 378,015 unique sgRNA combinations stably inserted in the D39V genome was created. In each cell, two sgRNAs are constitutively expressed along with an inducible dCas9, thus simultaneously targeting two different operons (Figure 1A). By challenging this library in standard growth conditions with and without a CRISPRi-activating inducer, 4,026 unique GIs were identified, involving both non-essential and essential genes. All interactions are made available and explorable through an online network browser (https://veeninglab.shinyapps.io/PneumoGIN). This work confirmed many previously identified GIs and revealed new connections between various essential processes. Specifically, we identified GIs between the putative membrane scramblase CozEa and the structural maintenance of chromosomes complex SMC and found several GIs between capsule formation and cell elongation. The methods and approaches described here can serve as a roadmap for dual CRISPRi-seq studies in other organisms.

**Figure 1.**
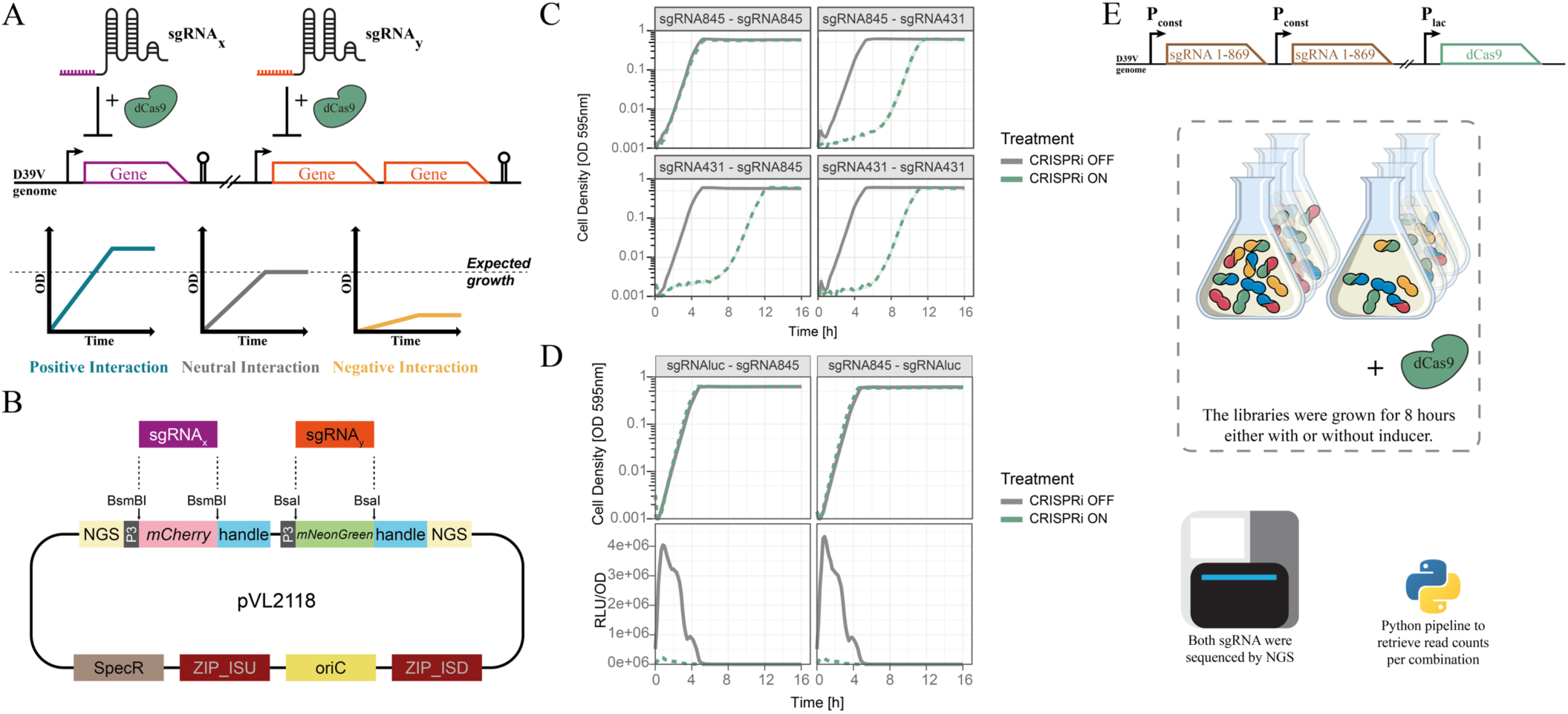
Workflow of dual CRISPRi-seq. **A.** Principles of the dual CRISPRi system. Two sgRNAs targeting two different operons containing one (purple) or more (orange) genes are co-expressed with dCas9, resulting in repression of two targets at the same time. Definitions of positive, negative, and neutral (or no) interactions are illustrated with schematics depicting logarithmic growth curves of dual CRISPRi strains. **B**. The pVL2118 vector was designed for cloning of dual sgRNAs. It features two sgRNA insertion sites flanked by BsmBI or BsaI recognition sites, respectively. The coding sequences of mCherry and mNeonGreen were inserted at the cloning sites for sgRNA1 and sgRNA2, respectively. Additionally, DNA sequences for Illumina sequencing, denoted as “NGS,” were added to both ends of the dual sgRNA sequence. This design allows for a one-step PCR during Illumina amplicon library construction. It replicates in *E. coli* but integrates at the non-essential ZIP locus in *S. pneumoniae*. See figure S1. **C**. Functionality test of dual CRISPRi with sgRNA845 (control sgRNA) and/or sgRNA431 (targeting the essential gene *tarI*). Cell density was measured at 595nm every 10 min for 16 hours. CRISPRi was activated with IPTG (green dashed lines). **D**. Functionality test of the dual CRISPRi system with sgRNA845 and sgRNA*luc* targeting the reporter firefly luciferase. Cell density at 595nm (top row) and luciferase activity (RLU/OD, bottom row) were measured every 10min for 16h. CRISPRi was activated with 1mM IPTG (green dashed lines). **E**. The structure of the dual CRISPRi library and workflow of CRISPRi-seq. The pooled 869 library was inserted successively twice into strain DCI23 (*lacI*, Plac-*dcas9*), creating an 869×869 pooled dual sgRNA library. The library was grown in the absence (CRISPRi OFF) or presence (CRISPRi ON) of 1 mM IPTG for 8 hours. After isolating the genomic DNA and ligating Illumina adapters by PCR, both sgRNAs were sequenced and custom scripts were used to retrieve read counts per sgRNA combination (see Methods).

## Results

### A dual sgRNA CRISPRi system in *S. pneumoniae*

To investigate GIs in *S. pneumoniae*, we constructed an integrative vector that allows for the simultaneous knockdown of two targeted operons (Figure 1B). The vector, pVL2118, has two strong constitutive synthetic P3^26^ promoters, each controlling the expression of either *mCherry* or *mNeonGreen*, surrounded by BsmBI or BsaI restriction sites and the sgRNA scaffolds (Figure 1B). This facilitates the efficient replacement of either reporter by a specific 20-bp sgRNA spacer with compatible overhangs by digestion/ligation. To verify the functionality of this dual CRISPRi system, we substituted *mCherry* and *mNeonGreen* on pVL2118 with a non-targeting sgRNA as control (sgRNA845) and/or sgRNA431 which targets the essential *tarI* gene, and transformed the plasmids into a strain containing an IPTG-inducible *dcas9* integrated into a neutral locus on the *S. pneumoniae* D39V chromosome (strain DCI23). Growth experiments showed that cloning sgRNA431 in either position on the vector led to efficient repression as the expected growth defect was observed, whereas the non-targeting sgRNA845 did not lead to any growth defect (Figure 1C). Additionally, we tested the repression activity by using an sgRNA targeting firefly luciferase (*luc*) which is constitutively expressed in strain VL997 (Table S4). When CRISPRi was induced by IPTG, we observed a decrease in luminescence regardless of the position of the sgRNA on the integrated vector (Figure 1D). Together, these results demonstrate that both sgRNA positions within the vector are functional for CRISPRi.

To assess genetic interactions on a genome-wide level, we used a previous sgRNA library design encompassing 869 sgRNAs, targeting all the high-confidence operons in the serotype 2 strain D39V^27^. The targeted 869 operons covered 1,586 out of 2,146 annotated genetic elements of the *S. pneumoniae* strain D39V (Table S1). By crossing this library with itself in 2 rounds of sgRNA cloning, we obtained a total of 378,015 unique sgRNA combinations (869*868/2+869=378,015) (Figure S1). We transformed the pooled dual sgRNA library into a host containing an IPTG-inducible *dcas9* on the chromosome (see Methods). The pooled IPTG-inducible dual CRISPRi library was grown for 8 hours at 37℃ in the commonly used C+Y medium in the presence or absence of 1 mM IPTG to allow for dCas9 expression and phenotypic differentiation of each sgRNA combination in a liquid-based competition assay (Figure 1E). Genomic DNA was extracted, the dual sgRNA sequences were amplified by PCR and sequenced in a paired-end fashion on Illumina platforms (Figure 1E, S1)^1^. The experiment was performed twice with three replicates each, and sgRNAs were sequenced on a NextSeq or NovaSeq sequencer (see Methods).

### Dual CRISPRi-Seq reliably identifies GIs

We filtered and kept high-quality reads (>Q30), obtaining 9.8M usable reads per sample on average (Figure 2A). To retrieve the read counts per sgRNA combination, we developed a python script derived from 2FAST2Q^28^ that allows simultaneous reading of both Read1 and Read2 fastq files, followed by sgRNA mapping and counting. Samples had an average of 24.25 and 27.79 reads per combination for non-induced and induced samples, respectively (Figure 2A). Out of the 378,015 unique possible combinations, only 9 sgRNA combinations were missing with 0 reads. 24,935 sgRNA combinations had an average read number of less than 10 in the non-induced samples (Table S2). This robust distribution underscores the thorough coverage of sgRNA combinations achieved. The induced samples displayed a distribution skewed towards 0 read counts, resulting from sgRNA combinations targeting essential gene pairs (Figure 2A). Principal component analysis (PCA) of the samples showed that the main source of variation between samples was CRISPRi induction, as 71.8% of variance was explained between non-induced and induced samples (Figure S2). In addition, pairwise Pearson correlation analysis revealed high similarity between induced samples (r > 0.8) (Figure S3).

**Figure 2.**
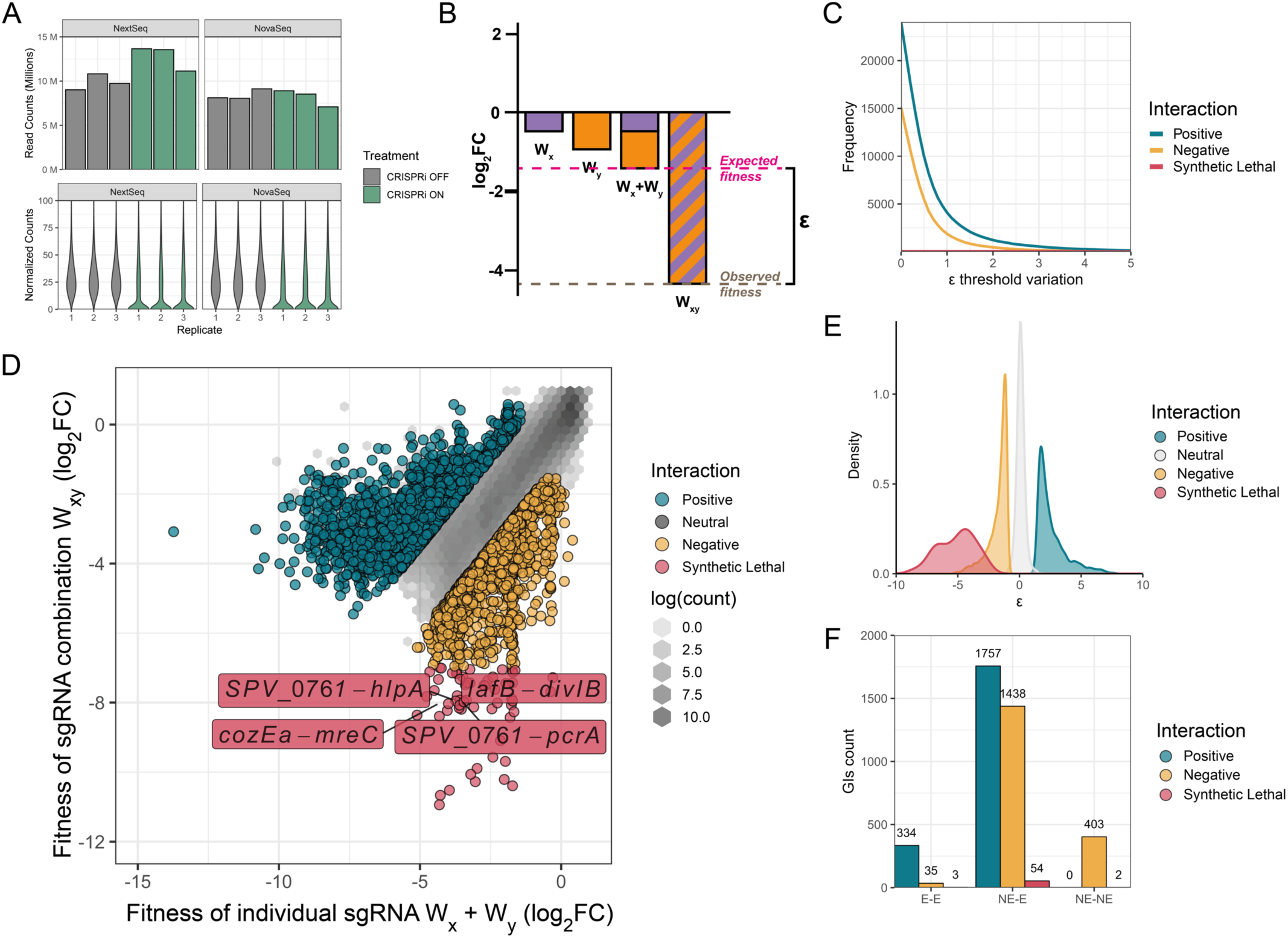
Fitness evaluation with the pooled dual sgRNA CRISPRi library. **A.** Bar plots showing normalized read counts in Million per sample (top row), before filtering. Violin plots showing the distribution of the normalized counts of each combination of sgRNA in each sample, before filtering (bottom row). The libraries were sequenced with NextSeq and NovaSeq6000. **B**. Schematic showing how epsilon (ε) is calculated, as described in Mani et al ^29^. W_x_ and W_y_ correspond to fitness of each individual sgRNA. W_xy_ is the fitness of the sgRNA combination. **C**. Variation of ε threshold when using the sum model ε = *Wxy* − *E*(*Wx* + *Wy*). As the threshold increases, the number of called GIs decreases. **D**. Identified positive, neutral, negative, and synthetic lethal genetic interactions using a threshold of 1.5 and 1 for positive and negative GIs, respectively. The hexagon shading shows the density of neutral points inside its area (natural log scale). Note that using these stringent thresholds, 4,026 significant GIs are called, representing approximately 1% of all possible GIs. **E**. Epsilon (ε) distributions of positive, neutral, negative and synthetical lethal combinations. **F**. Number of genetic interactions for essential-essential (E-E), non-essential-essential (NE-E) or non-essential-non-essential (NE-NE) sgRNA pairs.

We quantified the fitness effect of each sgRNA combination by enrichment analysis using the R package DESeq2 as described previously for CRISPRi-Seq^1^. For every sgRNA combination, we measured the depletion or enrichment in normalized read counts between non-induced and induced samples. The resulting log2 fold change (log2FC) directly quantifies the fitness of each sgRNA combination. As we have shown that the sgRNA position on the integrative vector has no influence on the targeted gene (Figure 1C-D), we summed read counts for the same combinations (e.g. sgRNA1-sgRNA2 or sgRNA2-sgRNA1). The individual fitness effect of each sgRNA is defined as the fitness of an sgRNA combination wherein the same sgRNA is present twice in the integrated vector (i.e. sgRNA1-sgRNA1). Note that using this double knockdown parameter per individual sgRNA may underestimate, but not overestimate, the strength of identified GIs with other sgRNAs. Using this definition, we calculated the strength of the GI described by epsilon (ε), by subtracting the observed fitness of two different combined sgRNAs (*Wxy*) with the expected fitness value *E*(*Wxy*), which we define as the sum of the individual fitness effects of each sgRNA (*Wx* and *Wy*), following the model ε = *Wxy* − *E*(*Wx* + *Wy*)(Figure 2B)^29^. The resulting ε score describes the deviation from the expected fitness loss of the sgRNA combination as a log2 fold change difference. To validate the ε model based on the sum of individual fitness scores, we compared it with ε distributions resulting from either minimum, multiplicative or logarithmic models as described before (Figure S4)^29^. As shown in Figure S4, the ε distribution resulting from the other models would make it very hard to properly segregate positive, negative, or synthetic lethal interactions. In addition, we verified the accuracy of the individual fitness effects of each sgRNA represented here by comparing *Wx* and *Wy* log2FC fitness values with our earlier published CRISPRi-Seq data that evaluated operon essentiality under similar growth conditions^25^. This gave a Pearson correlation of 0.96, indicating a robust justification for using *Wx* and *Wy* as individual fitness scores for each sgRNA (Figure S5).

To filter for high confidence GIs, we used an ε threshold in combination with the fitness scores *Wxy*, *Wx*, and *Wy*. We note that these threshold values are arbitrary and, depending on the research question, can be set either more or less stringent. We tested ε thresholds from 0 to 5, and determined that an ε threshold of -1 and 1.5 for negative and positive interactions, respectively, rejected most of the false positives and englobed already known GIs from the literature, such as the *mpgA-pbp2b* positive interaction^30^, and the *pbp1a-pbp2a* negative interaction (Figure 2C)^31,32^. Furthermore, we discarded sgRNA combinations when the number of reads of induced samples was below 4 and expected reads below 2 (see methods). Finally, we also filtered out observed positive interactions when two non-essential operons (p > 0.05, non-induced vs induced) were targeted, resulting from small fitness variations that are not significant. We defined a negative GI between two operons as an ε score smaller than -1 in combination with dual essentiality (*Wxy* p < 0.05). This signifies that the interaction between the two operons reduces overall fitness. Conversely, a positive GI is established when the ε score is greater than 1.5 (p > 0.05), reflecting a fitness increase. With these limits, we identified 4,026 interactions, including 1,935 negative GIs and 2,091 positive GIs, representing roughly 1% of all possible GIs. The majority of sgRNA combinations showed epsilon values close to zero, illustrating no difference between the expected fitness *E*(*Wxy*) and observed fitness *Wxy* (Figure 2D-E), and are hence identified as neutral interactions. To pinpoint putative synthetic lethal interactions, we set a combined fitness score of *Wxy* smaller than -7 reflecting a major fitness defect as stringent arbitrary threshold. This resulted in the identification of 59 putative synthetic lethal interactions including known synthetic lethal pairs of genes *cozEa* and *mreC* and genes *lafB* and *divIB* (Figure 2D)^33,34^. It also included gene *spv_0761* which is of unknown function to be synthetic lethal with *hlpA* and *pcrA* suggesting that *spv_0761* plays a role in chromosome segregation.

Interestingly, we identified more positive interactions than negative GIs when both targeted operons were essential (Figure 2F). This is in accordance with SGA data from *E. coli*, where essential-essential (E-E) pairs showed more positive than negative interactions^8,35^. This outcome is expected, given the challenge of identifying a negative interaction when the fitness loss, resulting from the presence of one sgRNA, is already severe. Finally, the ratio between negative and positive interactions was close to 1 for non-essential-essential pairs (NE-E)(Figure 2F).

### Revealing gene function via correlated GI analysis

To examine whether GIs are spatially enriched at specific regions of the chromosome, all identified GIs were mapped to the circular genome of D39V (Figure 3A, Figure S6). This showed no evident distribution pattern. However, when the number of GIs occurring along the genome were counted by dividing the genome into 50-kb segments, a high concentration of negative interactions between non-essential genes positioned between 0.5Mbp and 1Mbp of the D39V genome was observed (Figure 3B), suggesting some level of genetic redundancy specifically encoded on the right arm of the chromosome distal from the origin of replication.

**Figure 3.**
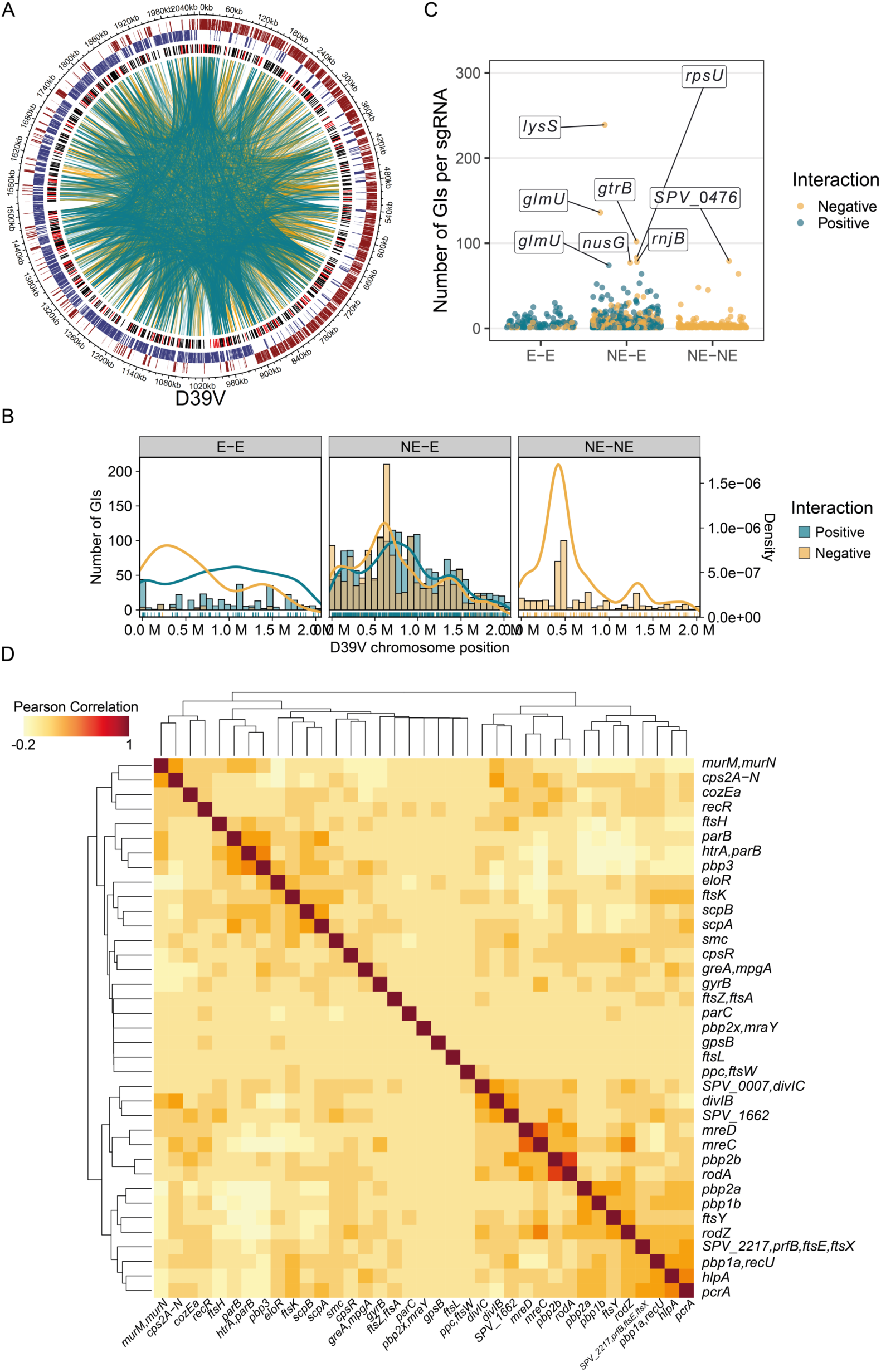
Visualization of genome-wide GIs identify highly connected genes and pathways. **A.** CIRCOS plot of identified significant GIs. From outside to inside: D39V chromosome position; Genes on the positive strand (dark red); Genes on the negative strand (dark blue); sgRNA targets, red represents essential sgRNAs, black represents non-essential sgRNAs based on Liu et al^25^. Negative (yellow curves) and positive (blue curves) interactions are shown as curves between two sgRNAs. **B**. Histogram and density plots showing the distribution of either negative (yellow) or positive (blue) interactions along the chromosome position, for essential-essential (E-E), nonessential-essential (NE-E) or nonessential-nonessential (NE-NE) sgRNA pairs. **C**. Scatter plot showing the sgRNA with number of GIs per sgRNA for essential-essential (E-E), nonessential-essential (NE-E) or nonessential-nonessential (NE-NE) sgRNA pairs. **D**. Clustered heatmap of Pearson correlations calculated across selected sgRNAs. The white (0) to dark red (1) gradient shows the Pearson correlation between each two sgRNAs.

The top 0.5% of sgRNAs ranked by their number of GIs, consist of interactions between non-essential and essential operons with the essential *lysS*, encoding for the lysyl-tRNA synthetase, having most GIs of all (NE-E, Figure 3C). A similar trend has been observed in *S. cerevisiae*, where essential genes tend to have more genetic interactions than non-essential genes^5^. These sgRNAs, all targeting essential genes, barely passed the here-applied filtering threshold, meaning that a significant growth defect is likely when combined with another sgRNA. Similarly, the same effect was observed by CRISPRi-TnSeq, where essential genes put cells on a threshold of collapse, which can be triggered by almost any other disturbance^15^. As sgRNAs targeting genes or pathways of similar function should have similar GI patterns, a Pearson correlation matrix of GIs was calculated across all sgRNA combinations, revealing sgRNA targets that are functionally closely related showing high correlation (Figure S7). We generated a smaller correlation heatmap by subsetting the correlation matrix and showing cell cycle related genes. This showed that genes involved in cell wall synthesis and cell division clustered together (*divIC*, *divIB*, *mreCD*, *pbp2B*, *rodA*), as well as genes involved in chromosome segregation and DNA damage (*ftsK*, *scpAB*, *smc*, *parB*) (Figure 3D). Importantly, a highly correlated GI is predictive for gene function and can thus be used for gene function prediction. On the basis of this, we propose that gene *spv_1662* is likely involved in cell division as it is highly correlated to GIs of genes *divIB* and *divIC*, known to be involved in late cell division (Fig. 3D).

To assess the function-relation distribution of genetic interactions, we categorized the targeted operons into Clusters of Orthologs Groups (COG) per gene. Some categories, such as cell wall biosynthesis and DNA replication, exhibited a greater prevalence of negative interactions over positive interactions (Figure S8A). A high positive to negative interaction ratio for the COG ‘defense mechanisms’ was observed, suggesting that refraining from investing in defense mechanisms might alleviate some of the stress induced by the repression of other operons under the tested experimental conditions (Figure S8B).

### Visualization of genome-wide GIs with PneumoGIN

To visualize the entire dataset, the identified GIs were mapped to a network, with each node representing a targeted operon or gene, and each edge an identified GI. Overall, the network highlights the connectivity between all processes in the cell. Unlike previous findings, which often demonstrated the clustering of functionally related genes in networks, our dual library approach draws attention to connections at the operon level. This is because most of the operons we studied already contain genes from the same pathways, such as the capsule operon or operons with ribosomal subunits^8,13^. To easily find specific GIs in this network, we developed PneumoGIN (<u>Pneumo</u>coccal <u>G</u>enetic Interaction <u>N</u>etwork), an online interactive R Shiny application. This application allows the user to search and explore the genome-wide GI network of the dual CRISPRi screening dataset. PneumoGIN is publicly available online (Figure 4) (https://veeninglab.shinyapps.io/PneumoGIN/)^27^.

**Figure 4.**
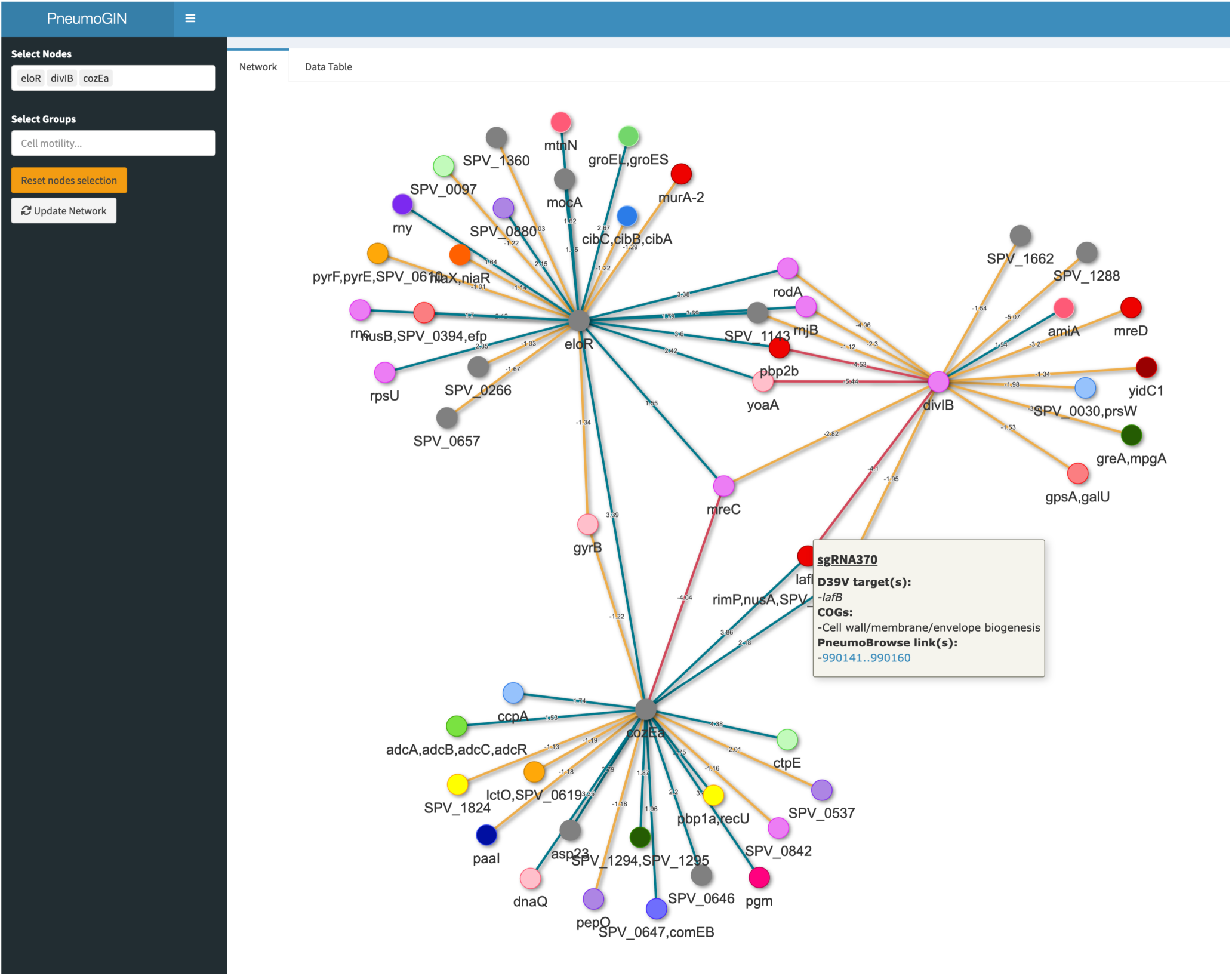
Snapshot of the PneumoGIN R Shiny App. The Pneumococcus Genetic Interaction Network (PneumoGIN) R Shiny app available at https://veeninglab.shinyapps.io/PneumoGIN/ allows users to select nodes of interest, and directly view predicted genetic interactions as a network involving the selected nodes. As an example, the *eloR*, *divIB* and *cozEa* genetic networks are shown. Positive, negative and synthetic lethal interactions are shown with blue, orange and dark red lines between nodes, respectively. Node colors are grouped relative to the predicted Clusters of Orthologous Groups (COG) of the targeted operon/gene.

### Dual CRISPRi-Seq is flexible and scalable

To show the versatility of the dual CRISPRi-Seq system, as well as its reproducibility, a smaller pooled library was constructed, by selecting 18 specific sgRNAs that were combined with our previously described 1,499 sgRNA pooled library, targeting 2,111 out of 2,146 genetic elements in the D39V genome ^25^. The 18 sgRNA’s were chosen to include genes we have worked on in the past such as genes involved in chromosome segregation (*smc*), DNA replication (*ccrZ*) and cell division (*ftsZ*), or genes that had interesting interaction partners as observed in the genome-wide dual CRISPRi-Seq screen such as *cozEa*. Along with the 18 selected sgRNAs, a 19^th^ sgRNA, targeting the firefly luciferase gene (*luc*) was introduced. This *luc*-targeting sgRNA was used as a control for fitness calculations of each combination as the pneumococcal genetic background used for library construction did not contain *luc*. The 19×1,499 dual library was constructed using a process similar to that of the 869×869 dual library (Figure S1). This resulted in a “mini” library of 26’964 unique combinations (18*1,499-18), assessing the genetic interactions of the 18 selected target operons on a complete genome-wide level. After induction and sequencing of the library, the fitness scores of each of the 26’964 combinations were compared with the fitness score of each sgRNA combined with the sgRNA targeting *luc*. Interestingly, we found that genes *spv_1836* and *spv_1482*, both of unknown function and not present in the 869×869 library, had strong negative GIs with *rodA*, suggesting they play a role in cell elongation (Table S7).

By comparing GIs between the mini library and the genome-wide library, we observed a strong, positive Pearson correlation for sgRNA pairs that had a significant interaction in both (R = 0.85, p < 0.001, Figure S9), and a moderate positive correlation when the interaction was significant in at least one library (R = 0.38, p < 0.001, Figure S9). This highlights how a custom dual sgRNA library can be set up to either reproduce or expand previous results, as well as showing the scalability without losing sensitivity of the here-described dual CRISPRi-Seq system.

### Validation of the dual CRISPRi-Seq GI network

To validate the dual CRISPRi-Seq data, we selected 23 predicted GIs for further analysis on the basis of including genes spanning diverse essential processes in the cell such as cell elongation, cell division, chromosome segregation and central metabolism (Figure S10). We individually constructed 23 double CRISPRi strains, replacing *mCherry* and *mNeonGreen* from pVL1998 with one or two selected sgRNAs (Figure 5A). Next, fitness was assessed by measuring growth over 16 hours, and the area under the curve (AUC) was calculated over the first 8 hours. To be able to compare with the epsilon ()) values obtained from the genome-wide dual CRISPRi-Seq screen, we measured the difference between the growth fitness and the expected growth fitness in each dual CRISPRi strain using a multiplicative model, defined by Δ*W_AUC_* = *W_AUCxy_* − *E*(*W_AUCx_* * *W_AUCy_*) (See Methods)^15^. The Δ*W_AUC_* and ε values showed a strong positive Pearson correlation (R = 0.73, p < 0.001, Figure 5B), reflecting the robustness of the large pooled dual screen and reproducibility of the results with individual CRISPRi strains. Specifically, we validated previously reported positive genetic interactions using dual CRISPRi, including *rodA-eloR, pbp2b-mpgA*, and negative interactions such as *pbp1a-pbp2a* (Figure 5C, Figure S10)^30^. In total, 21 out of the 23 constructed dual CRISPRi strains demonstrated the expected growth pattern (Figure S10).

**Figure 5.**
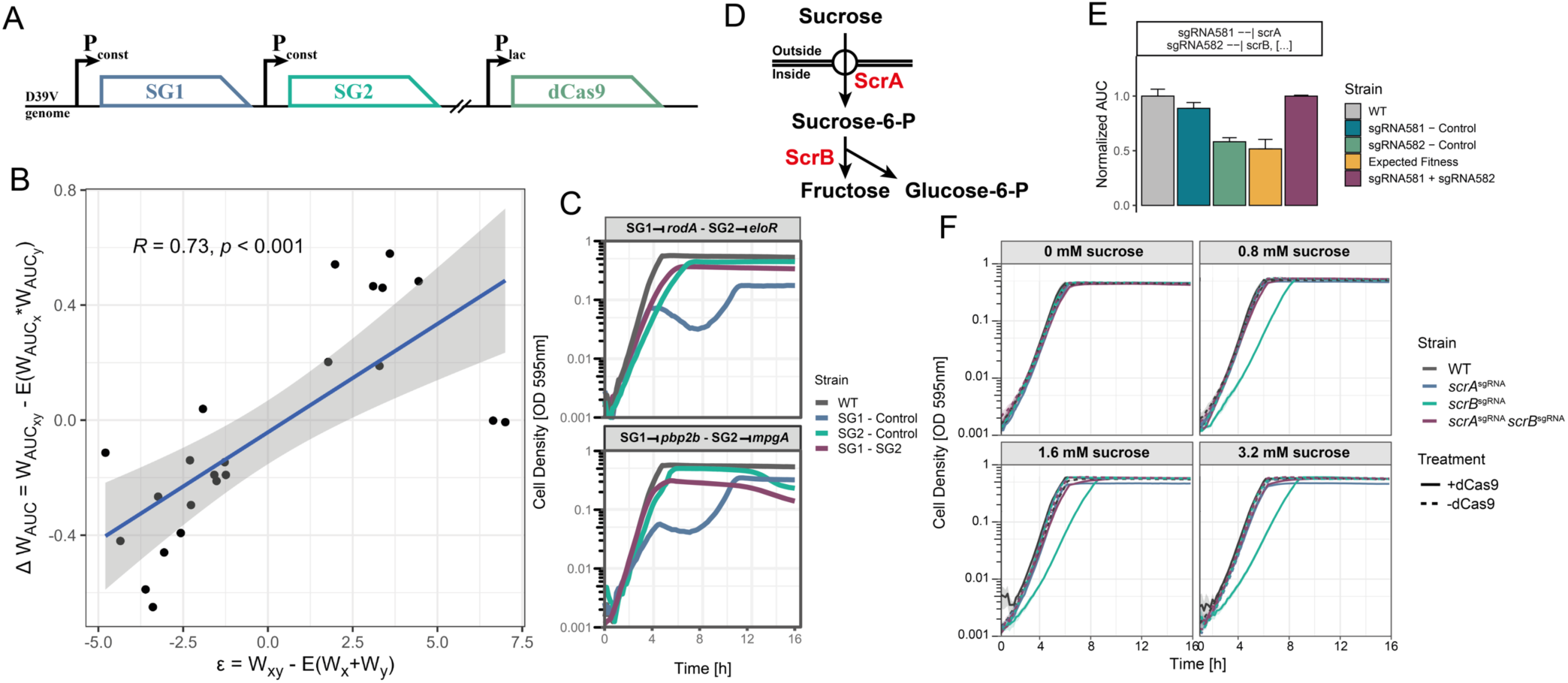
Validation of GIs identified by dual CRISPRi-Seq. **A.** Selected sgRNAs were either cloned in position SG1 or SG2 and integrated into the D39V genome at the ZIP locus^44^. IPTG-inducible *dcas9* is integrated at the *bgaA* locus^9^. **B**. Validation of 23 selected sgRNA combinations by growth curves. Optical density at 595nm was measured every 10 min. The area under the growth curve (AUC) was calculated for the first 8 hours and DW_AUC_ was calculated as described in the methods. The DW_AUC_ value for each sgRNA combination was compared with the epsilon values of the sgRNA combination evaluated by dual CRISPRi-Seq. Linear regression (blue line) with confidence interval in grey, and corresponding Pearson correlation coefficient R along with the p-value are shown. **C**. Growth curves of two dual CRISPRi strains. Only growth curves when CRISPRi was activated by IPTG are shown. Purple lines show the strain where both SG1 and SG2 are present. Blue lines when only SG1 is present. Green lines when only SG2 is present. Top facet shows the positive interaction between *rodA* and *eloR*, and bottom facet shows positive interaction between *pbp2b* and *mpgA*. **D**. Metabolic pathway showing the uptake of sucrose by ScrA, followed by the hydrolysis into fructose and glucose-6-P by ScrB. **E.** Normalized area under the curve (AUC) of the first 8 hours of growth of sgRNA581 and sgRNA582, targeting *scrA* and *scrB* respectively. The yellow bar shows the expected fitness between sgRNA581 only (blue bar) and sgRNA582 only (green bar) (see methods). The purple bar indicates the measured fitness when both sgRNA581 and sgRNA582 are present in the strain. **F**. Optical density measured every 10 min for 16h with different concentrations of extracellular sucrose (0 mM, 0.8 mM, 1.6 mM, and 3.2 mM). The lines represent different conditions: no sgRNA present (grey), only sgRNA581 targeting *scrA* (blue), only sgRNA582 targeting *scrB* (green), and both sgRNA581 and sgRNA582 present (purple). The dashed line represents CRISPRi being off, while the solid line represents CRISPRi being activated by the presence of IPTG.

Looking more precisely at some of the previously unknown identified genetic interactions, sgRNA581 targeting the sucrose phosphotransferase transporter gene *scrA* and sgRNA582 targeting the sucrose-6-P hydrolase *scrB* were predicted to have a positive interaction (Figure 5D). Indeed, dual CRISPRi targeting *scrA* and *scrB* showed a significant Δ*W_AUC_* between the expected fitness and the observed fitness of the combination, showing that the double knockdown masks the expected fitness loss (Figure 5E). Based on the predicted function of ScrA and ScrB, we investigated the presence of different concentrations of sucrose in the growth medium on this positive interaction. When sucrose was not present in the medium, no growth defect was observed whether *scrA* and/or *scrB* genes were repressed by CRISPRi. However, when sucrose was present in the medium, a reduced growth rate was observed only when *scrB* transcription was repressed. However, this reduced growth rate was restored when both *scrA* and *scrB* were knockdown simultaneously by dual CRISPRi (Figure 5F). This result suggests that the growth defect resulting from the knockdown of *scrB* is contingent upon the presence of sucrose. ScrB is a sucrose-6-P hydrolase which can transform sucrose-P into fructose and glucose-6-P ^36^. Therefore, we hypothesized that in the *scrB* knockdown strain, there is an increased intracellular accumulation of sucrose-P by ScrA, which subsequently impedes the growth of *S. pneumoniae*. Indeed, it has been shown before that phosphorylated sugar metabolites can be toxic upon their intracellular accumulation^37^.

### SMC interacts with genes involved in chromosome segregation and division

The pneumococcal GI network reveals many interactions among genes involved in chromosome segregation and division. Of these genes, *smc*, encoding the structural maintenance of chromosome protein, was previously identified as non-essential although it was found to be important for pneumococcal chromosome segregation^38^. As *smc-scpA-scpB* forms a triad in *Bacillus subtilis* to form bacterial condensin, or the SMC complex^39^, it is expected to find them interacting closely in the network. Strikingly, we found key genes known to be involved in chromosome segregation to negatively interact with one or more members of the SMC triad, such as *hlpA* encoding the histone like protein HlpA (HU), and *ftsK*, encoding the conserved ATP-dependent DNA translocase (Figure 6A)^40^. Although both *hlpA* and *ftsK* are already essential by themselves, the negative interactions with *scpA* and/or *scpB* further illustrate the importance of functional redundancy. This highlights the strength of dual CRISPRi in that genetic interactions between essential genes can be quantified.

**Figure 6.**
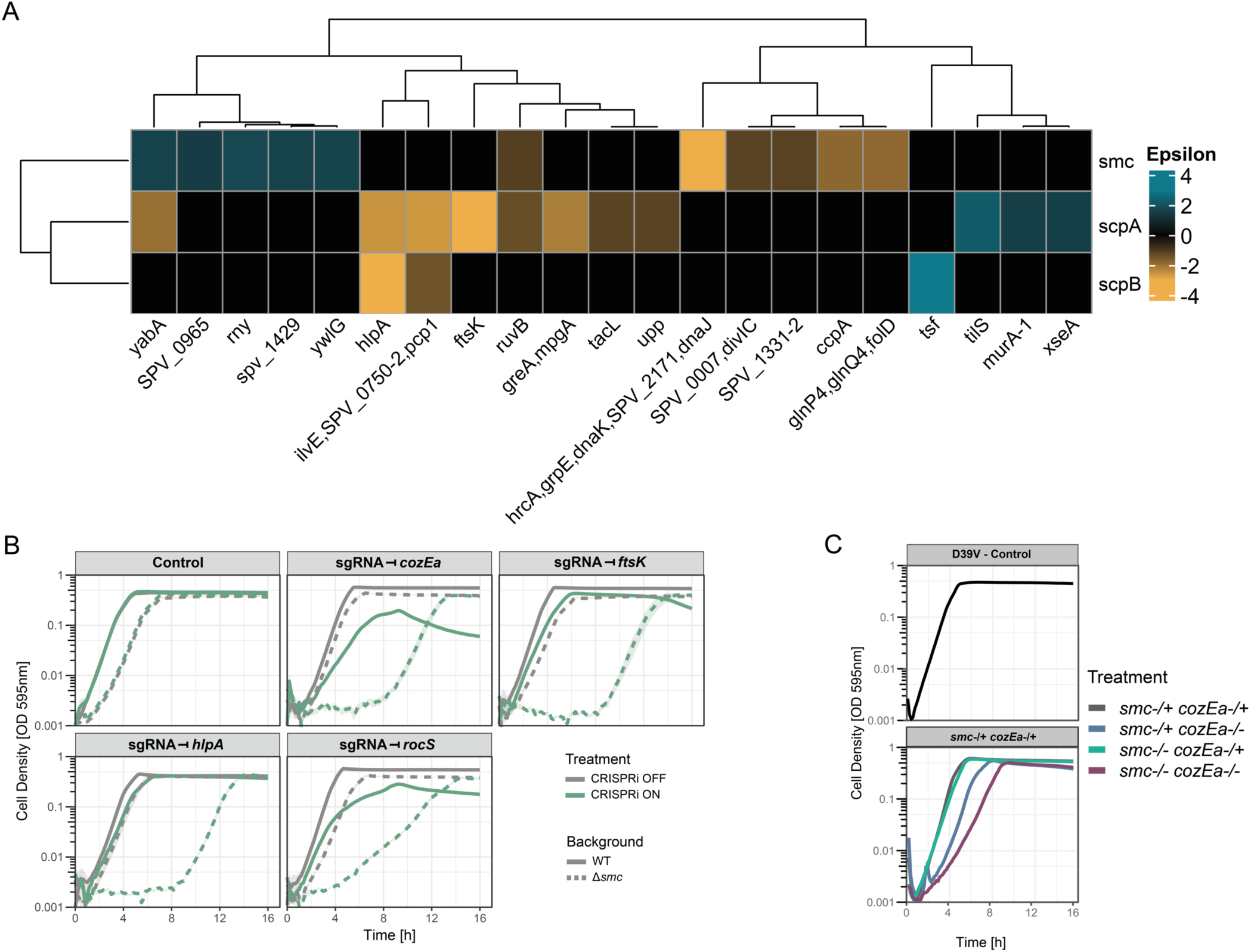
Genetic interactions within the chromosome segregation pathway. **A.** Clustered heatmap of all significant genetic interactions involving *smc*, *scpA* or *scpB*. The blue-to-yellow gradient shows the strength of positive to negative genetic interaction. **B**. Confirmation of hits interacting with *smc* by CRISPRi. s*mc* deletion was inserted in different CRISPRi strains with sgRNA targeting either *cozEa, ftsK, hlpA, rocS*. Growth was assessed for 16 hours, and OD was measured at 595nm every 10 minutes. **C**. Double depletion of *smc* and *cozEa* by ectopic expression via the IPTG-inducible promoter Plac for *smc*, or the aTc-inducible promoter for *cozEa* (VL4072, Table S4). Growth was assessed for 16 hours, and OD was measured at 595nm every 10 minutes.

To verify genetic interactions involving *smc*, we inserted individual sgRNAs targeting the known chromosome biology genes *ftsK*, *rocS* and *hlpA* into an *smc* deletion mutant, thus avoiding any unexpected polar effects. Consistent with the dual CRISPRi-Seq findings, all combinations showed large growth defects when the targeted gene was knocked down by CRISPRi in the *smc* deletion mutant (Figure 6B). We also examined a putative and surprising interaction between *cozEa*, which is involved may be a membrane scramblase involved in midcell localization of *pbp1a* and interacts with *pbp2x*^33,41–43^, and *smc*. While this GI fell just below our stringent cutoff criteria, knocking *cozEa* down by CRISPRi led to a synthetic lethal phenotype in the *smc* mutant (Figure 6B).

To further validate the GI between *cozEa* and *smc*, in-frame double deletion mutants were made, which were complemented by inserting an ectopic copy of each operon under either an IPTG inducible promoter (Plac) at the non-essential ZIP locus or anhydrotetracycline (aTc) inducible promoter (Ptet) at the *bgaA* locus^44^. As shown in Figure 6C, growth of double depleted *smc*/*cozEa* cells was hampered compared to single depleted cells. Single cell analysis of double depleted cells showed that cells were elongated after gene depletion, highlighting the important interaction and balance between cell division and chromosome segregation (Figure S11).

Overall, positive interactions with the SMC triad complex tend to include genes that putatively reduce the speed of the cell cycle to allow more time for the cell to organize and segregate the chromosome. This agrees with the fact that *smc* mutants in *B. subtilis* are only viable under slow growth conditions or in the presence of sublethal concentrations of antibiotics that lead to reduced replication fork velocity^45^. On the contrary, genes negatively interacting with the SMC triad will tend to increase stress around chromosome organization or give less time for the cell to properly segregate its chromosome (such as *hlpA*, *ftsK*, or *rocS*)^46^. The information provided by the above-mentioned function-known genes could also shed light on the function of many hypothetical genes found in the same interaction network, including *spv_0965, spv_1429, spv_1331-2, spv_0750-2* (Figure 6A). These genes were identified by dual CRISPRi-Seq to genetically interact with members of SMC triad, suggesting their involvement in cell division and chromosome segregation, and are prime candidates for follow-up studies.

### DivIB and DivIC at the center of a cell elongation fail-safe mechanism

In the oval-shaped pneumococcus, cell division and cell elongation are two separate processes that rely heavily on septal and peripheral peptidoglycan synthesis^47^. To better understand how these two processes are coordinated to enable proper cell morphology, we investigated GIs between division and elongation genes. An interesting trend that is observed in the dual CRISPRi network are GIs involving DivIB and DivIC. The two proteins were reported to form a late-divisional complex with FtsL at the septum and thus be involved in the same pathway^47–49^. Furthermore, DivIC was reported to be involved in the recruitment of the septal peptidoglycan synthase complex FtsW-PBP2x^49^. It was proposed that in *Pseudomonas aeruginosa*, DivIB is in complex with DivIC and FtsL, involved in the regulation of septal PG synthases, such as FtsW^47,50^. Consistent with the tightly related function between DivIB and DivIC, pneumococcal *divIB* and *divIC* shared many negatively interacting genes such as *rodA, mreC, mreD, pbp2b,* and *mpgA*, whose functions are primarily part of the elongation machinery (Figure 7A)^30,47,48^. These findings suggest that when septal peptidoglycan synthesis is impaired by knocking down *divIB* or *divIC*, fine-tuning of the elongasome by *rodA, mreC, mreD, pbp2b,* and *mpgA* becomes more vital. The GI network of *divIB* and *divIC* also includes uncharacterized genes, such as *spv_1662* and *spv_1143* (Figure 7A). Separately cloned dual CRISPRi confirmed the observed negative interaction in dual CRISPRi-seq between *spv_1662* and *mreD* (Figure 7B). Together, these data strongly suggest that these uncharacterized genes have a role in cell division, cell elongation or cell envelope homeostasis.

**Figure 7.**
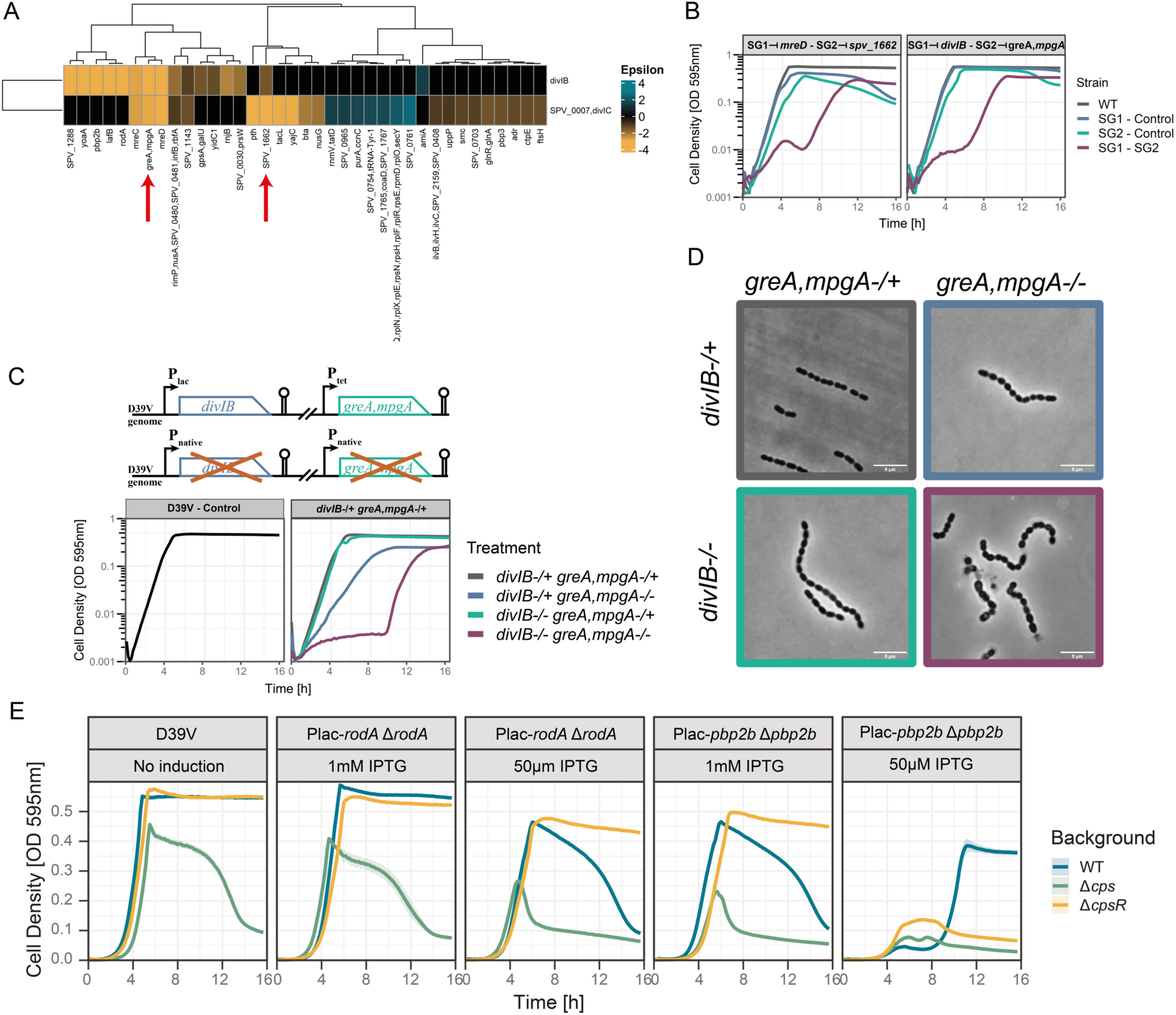
Genetic interactions among genes involved in cell elongation and capsule biosynthesis. **A.** Clustered heatmap showing GIs involving the *divIB* or *spv_0007*-*divIC* operon. Note that *spv_0007* and *divIC* are in the same operon and targeted jointly by one sgRNA. The blue-to-yellow gradient shows the strength of positive to negative GIs. Red arrows highlight the key genes described in text. **B**. Growth of dual CRISPRi strains, optical density at 595nm was measured every 10 minutes for 16h. Only growth curves with activated CRISPRi are shown. Purple line shows the strain where both SG1 and SG2 are present. Blue line when only SG1 is present. Green line when only SG2 is present. **C**. Double depletion of *divIB* and *greA-mpgA* operon by ectopic expression via the IPTG-inducible promoter Plac for *divIB*, or the aTc-inducible promoter for *greA-mpgA* operon (VL4072, Table S4). Growth was assessed for 16 hours, and OD was measured at 595nm every 10 minutes. Grey line indicates the presence of both IPTG and aTc. Blue line represents the presence of only IPTG. Green line indicates the presence of only aTc. Purple line indicates the absence of either inducer. **D**. Phase contrast microscopy of the double depletion of the *divIB* and *greA-mpgA* operons. Colours match the conditions described in C. **E**. Complementation/deletion of either *rodA* or *pbp2b* was constructed in WT (blue line), D*cps* (green line, *cps2A-O* deletion) or D*cpsR* (yellow line) strains. Growth was assessed for 16 hours, and OD was measured at 595nm every 10 minutes. For clarity, cell density is plotted on a linear scale.

Another interesting negative genetic interaction was observed between *divIB* and the muramidase *mpgA* (also known as *mltG*)^30^, which was confirmed by dual CRISPRi and by in-frame double deletion/complementation assays (Figures 7B, 7C, S10). The phenotype was also confirmed by microscopy, and the depletion of both operons led to round cells with heterogenous sizes (Figure 7D). Overall, the genetic interactions involving *divIB* and *divIC* highlight the significant roles of these genes in the pneumococcus, despite being non-essential. This implies that the pneumococcus employs redundancy mechanisms to ensure proper elongation, division, and growth.

### The capsule has a structural function supporting failures of the elongasome

The capsule is a major virulence factor of *S. pneumoniae* and can at the same time mask cell division defects in certain genetic backgrounds^51,52^. However, the full extent of phenotypes that are masked by the capsule is unknown, highlighting the importance of understanding variations of gene essentialities in the presence and absence of the capsule. Dual CRISPRi-Seq identified negative genetic interactions between the capsule operon (*cps2A-N*) and both *rodA* and *pbp2b*, which are involved in cell elongation (Figure S12, Table S3). Consistently, positive interactions between the capsule operon repressor *cpsR* with *rodA* and *pbp2B* were also identified (Figure S12, Table S3). To confirm these positive and negative interactions involving the capsule and genes involved in cell elongation, we deleted either the capsule operon or its negative regulator in either a Plac-*pbp2b* Δ1*pbp2b* strain or a Plac-*rodA* Δ1*rodA* (*rodA*-/+) strain. An increased lysis of the *rodA-/+* mutant in the capsule operon deletion background was observed when 50 μM IPTG was added, which explained the identified negative genetic interactions between the capsule operon and *rodA* (Figure 7E). In the *cpsR* mutant background, *S. pneumoniae* survived relatively better at 50 μM IPTG than the wild-type control (Figure 7E), confirming the positive interaction between *rodA* and *cpsR*. Comparing the essentiality of *pbp2b* in a *cps* mutant with the wild-type strain, we observed a decreased fitness at 50 μM IPTG in the *cps* mutant, but a higher fitness when IPTG was present (Figure 7E). When *cpsR* was deleted, lysis at stationary phase was lower than the wild type background when 50 μM IPTG was present, suggesting *cspR* deletion improves cell integrity when *pbp2b* is depleted. This confirms the positive interaction between *pbp2b* and *cpsR*. Together, these genetic interactions involving the capsule emphasize the importance of the capsule for cell integrity. They show that cell lysis caused by reduced peripheral peptidoglycan synthesis can be counteracted by the presence of a capsule, which thus functions to structurally reinforce the cell. These data predict that unencapsulated bacteria are more susceptible towards cell wall targeting antibiotics that preferentially target PBPs involved in elongation such as piperacillin, which has a relatively high affinity for *S. pneumoniae* Pbp2B^53^.

## Discussion

Here we describe dual CRISPRi-Seq to explore GIs on a genome-wide scale in *S. pneumoniae*. Compared with previous genome-wide screening methods to map GIs such as Tn-Seq, or more recently CRISPRi-TnSeq^7,8,10,12,15,54^, dual CRISPRi-Seq allows the study of GIs involving both essential and non-essential genes in an unbiased way. An integrative vector that can accept two sgRNAs by subsequent digestion/ligation was created giving the possibility to insert any kind of library, as well as changing homologous regions that would allow the integration into another species’ chromosome. However, the library size and/or the chosen organism should allow for the recovery of enough transformants to ensure all sgRNA combinations are present within the constructed pool. Using this genetic platform, a pooled library containing 378,015 unique combinations of sgRNAs covering over 70% of all possible pairwise genetic interactions of the *S. pneumoniae* D39V genome was generated. As a proof of concept, we grew the library in rich medium, with or without CRISPRi activation. The resulting enrichment of sgRNA pairs was determined by paired-end sequencing. By developing a bioinformatic pipeline for dual sgRNA data, 1,935 negative and 2,091 positive high confidence GIs were identified, most of which have never been described. The generated network of GI can be visually explored through PneumoGIN, a publicly accessible online R shiny app (https://veeninglab.shinyapps.io/PneumoGIN)(Figure 4). While we cannot delve into every uncovered genetic interaction due to the vastness of the network, we emphasized major patterns that illustrate the use of dual CRISPRi-Seq to identify and investigate genetic interactions.

Previous work used temperature sensitive (TS) mutant alleles to query genetic interaction of essential *S. cerevisae* genes^3,5^, but the generation of TS alleles is cumbersome and not possible for all genes within a genome, nor for many species. In *E. coli*, a set of hypomorphic mutants was used to assess genetic interactions of a set of selected essential genes^7^. Similar to these *S. cerevisiae* genetic interaction studies, we also observed a tendency where essential genes had more GIs than non-essential genes^3,5^. This was also observed in *E. coli*, using SGA to screen for GIs, where essential genes were found to be more highly connected in the network compared to non-essential genes^7^. Compared with SGA or Tn-Seq, our CRISPRi approach may result in unwanted polar effects. We designed our library based on high confidence TSS of the D39V genome to cover the maximum number of genes with a small library, thereby limiting the coverage needed for all combinations to be present during the construction of the library. A future improvement of the dual CRISPRi library would be to increase the library size to have at least one sgRNA per open reading frame in the D39V genome. However, such an increase in library size would also require harvesting more colonies and deeper sequencing to preserve coverage, as the number of unique combinations would be higher.

Validation experiments demonstrated a high confirmation rate, accentuating the robustness and reliability of this method. Compared to our recent study employing the combination of CRISPRi and Tn-Seq to investigate GIs in *S. pneumoniae*, the obtained GI profiles of specific genes had overlap, such as with the negative interactions involving *smc-scpA-scpB* and *cozEa*^15^. We confirmed strong negative interactions with the SMC triad, such as HlpA, FtsK and RocS. As *S. pneumoniae* does not have a nucleoid occlusion system as in *B. subtilis* or *S. aureus*^7,55–57^, these GIs involving the SMC triad reveal the importance of a proper balance in the various partners that play a role in chromosome segregation. Correlation analysis identified several as of yet uncharacterized genes that may be involved in chromosome segregation which are prime candidates for follow-up studies. We further demonstrated the importance of cell elongation when cell division is impaired. Specifically, when DivIC or DivIB are absent, the requirement of many cell elongation proteins is augmented showcasing the interconnected relationships that exist genome-wide between distinct processes. In addition, we identified uncharacterized genes such as *spv_2029, spv_1662 and spv_1143* that negatively interacted with some of the cell elongation and cell division members. This provides prospects in understanding their function and their involvement in the cell cycle, and possibly forge the way into the understanding of new mechanisms that could be used for drug development. Finally, the here-described dual CRISPRi-Seq platform sets the stage for more complex conditions, such as the presence of antibiotics to uncover gene-gene-drug interactions and different growth conditions that would mimic infection-relevant conditions^58^. This would lead to the identification of genetic interactions that are medium-, or antibiotic-specific, potentially revealing possible therapeutic avenues for combinatorial therapies. The here-described methods and bioinformatic approaches can serve as a roadmap for genome-wide GI studies in other organisms and the generated pneumococcal GI network can serve as a starting point for new biological discovery and translational research.

## Methods

### Bacterial strains, growth, and transformation

Bacterial strains used in this project are a derivative of the serotype 2 *S. pneumoniae* D39V^27^. *S. pneumoniae* was grown in C+Y medium at pH 6.8, adapted from de Bakker et al^1^. Plasmids, ligation products or genomic DNA were transformed into *S. pneumoniae* cells after inducing with competence stimulation peptide 1 (CSP-1) as previously described^59^. When required, induction of the aTc-inducible promoter (Ptet) was carried out by supplementing the medium with 50 ng/mL aTc (anhydrotetracycline, Sigma-Aldrich) and the IPTG inducible promoter (Plac) was activated with 100 μM IPTG (β-D-1-thiogalactopyranoside, Sigma-Aldrich). Transformants were selected on Columbia agar with 2% sheep blood at 37°C with 5% CO2 using appropriate antibiotics concentration (4.5μg/ml chloramphenicol, 0.5μg/ml erythromycin, 250μg/ml kanamycin, 100μg/ml spectinomycin, 10μg/ml trimethoprim). Inserted constructs were checked by PCR followed by sanger sequencing at Microsynth AG. All strains were stocked at OD_595nm_ 0.3 at -80°C with 15% glycerol.

All strains listed in Table S4 were constructed via integration of a linear DNA fragment into the chromosome by homologous recombination. The linear DNA normally possesses an upstream and downstream homology region of ∼1kb, and an insert, assembled using standard Golden Gate Assembly. Either Esp3I, BsaI, or AarI sites were directly integrated into the primers. All integrated constructs were confirmed by PCR and the resulting fragment sequenced. For gene deletions, an antibiotic cassette (either erythromycin, kanamycin, trimethoprim), was used for selection, and an in-frame replacement of the open reading frame by the resistance marker was constructed, to reduce any potential polar effect. For cloning an ectopic copy of the gene under a P_tet_ promoter, a tetracycline resistance marker was used, and the resulting fragment was integrated into the *bgaA* locus, as described before^25^. For cloning an ectopic copy of the gene under a P_lac_ promoter, a pPEPZ vector was used, containing spectinomycin resistance marker, and the resulting assembly was integrated into either the *zip* locus or *cep* locus, as described before^44^. Note that the ectopic copy of the gene was always inserted before the deletion of the original loci, to prevent unwanted suppressor mutations.

For VL4068 (Table S4), VL333 was used as the parent strain. For *zip::*P_lac_*-divIB,* OVL2195 and OVL2196 were used to amplify *divIB* from genomic DN (gDNA) A of D39V. OVL2181 and OVL2182 were used to amplify pPEPZ-spc-P_lac_ vector. For *divIB::ery* construct, OVL2344 and OVL2345 were used to amplify the upstream region of *divIB*. OVL2346 and OVL2347 were used to amplify the downstream region of *divIB*. OVL4823 and OVL1783 were used to amplify erythromycin resistant cassette. For *bgaA*::P_tet_-*mltG_greA* construct, OVL5024 and OVL5025 were used to amplify *mltG_greA* operon. OVL2077-OVL5018, and OVL5019-OVL1369 were used to amplify the *bgaA* upstream-*tetracycline*-P_tet_ and *bgaA* downstream, respectively. For *mltG_greA::kan*^R^, OVL5336-OVL5337 and OVL5338-OVL5339 were used to amplify *mltG_greA* upstream and *mltG_greA* downstream, respectively. OVL3981 and OVL3982 were used to amplify kanamycin resistant cassette. Each construct was sequentially transformed.

For V4072 (Table S4), VL333 was used as a parent strain. For *zip::*P_lac_*-smc,* OVL4793 and OVL4794 were used to amplify *smc* from D39V gDNA. OVL2181 and OVL2182 were used to amplify pPEPZ-spc-P_lac_ vector. For *smc::trmpR* construct, OVL2509 and OVL2510 were used to amplify *smc::trmp*^R^ from VL404 (Van Raaphorst, R. *et al*, 2017). For *bgaA*::P_tet_-*cozEa* construct, OVL5036 and OVL5037 were used to amplify *mltG_greA* operon. OVL2077-OVL5018, and OVL5019-OVL1369 were used to amplify the *bgaA* upstream-*tetracycline*-P_tet_ and *bgaA* downstream, respectively. For cozEa*::ery*^R^, gDNA of VL91 was used^9^.

For VL2477 (Table S4), VL236 was used as parent strain. For *zip::*P_lac_*-rodA,* OVL2191 and OVL2192 were used to amplify *rodA* from D39V gDNA. OVL2181 and OVL2182 were used to amplify pPEPZ-spc-P_lac_ vector. For *rodA::ery* construct, OVL2348 and OVL2349 were used to amplify the upstream region of *rodA*. OVL2350 and OVL2351 were used to amplify the downstream region of *rodA*. OVL4823 and OVL1783 were used to amplify erythromycin resistant cassette.

For VL6797, and VL6798, VL2477 or VL3924 were used as parent strain. *cps*::*chl*^R^ was amplified with OVL4609 and OVL4611 from VL4146. For VL6799 and VL6800, VL2477 or VL3924 were used as parent strain, respectively. For *cpsR*::*kan*^R^ construct, OVL9610 and OVL9611 were used to amplify the upstream region of *cpsR.* OVL9612 and OVL9613 were used to amplify the downstream region of *cpsR*. OVL9608 and OVL9609 were used to amplify the kanamycin resistance cassette.

For VL6792, VL6793, VL6794, and VL6795, VL2900 was used as the parent strain (Table S4). The individual sgRNA was cloned into the pPEPZ-sgRNAclone plasmid in *E. coli*, as described previously^20^. The integrative plasmid was then transformed into VL2900.

### Construction of the pPEPZ-P3-dualsgRNA vector

This vector was constructed based on vector pPEPZ-sgRNAclone (addgene #141090)^20^. The DNA fragment with P3-BsaI-mNeonGreen-BsaI was ordered as a gBlock and then cloned into pPEPZ-sgRNAclone by infusion cloning. The P3-BsaI-mNeonGreen-BsaI gBlock was amplified with two primers (For: CCATTCTACAGTTTATTCTTGACATTG and Rev: GGATCCATGAGTTTTTGTTCGGG). The pPEPZ-sgRNAclone

vector was linearized for infusion cloning by PCR with two primers:

- For: TAAACTGTAGAATGGCTGTCTCTTATACACATCTGACG
- Rev: AAAACTCATGGATCCCCATTCTACAGTTTATTCTTGACAT

The infusion reaction was performed with the amplified gBlock and linearized vector in accordance with the manufacturer’s instructions (Vazyme, cat. C113-02). The infusion reaction was transformed into the *E. coli* Stbl3 competent cells. The sequence of the gBlock is provided below:

P3-BsaI-mNeonGreen-BsaI gBlock, 5’-3’:

CCATTCTACAGTTTATTCTTGACATTGCACTGTCCCCCTGGTATAATAACTATATGAGACctAGGAGGACAAACATGTCAAAAGGAGAAGAGCTGTTCACAGGTGTTGTGCCGATTCTCGTTGAGCTTGACGGAGATGTAAACGGACACAAATTCTCTGTTCGCGGTGAAGGTGAAGGAGATGCAACAAACGGCAAGCTGACATTGAAGTTTATTTGCACAACTGGAAAGCTGCCGGTTCCTTGGCCGACACTTGTAACGACGCTGACTTACGGCGTTCAATGCTTCTCTCGTTATCCAGACCACATGAAACGCCATGATTTCTTCAAATCTGCAATGCCTGAAGGCTACGTTCAAGAGCGTACGATCAGCTTCAAAGATGACGGAACGTACAAAACAAGAGCAGAAGTGAAGTTTGAAGGTGACACACTTGTGAACCGCATTGAATTGAAAGGCATTGATTTCAAAGAAGATGGAAACATCCTTGGACACAAACTTGAATACAACTTCAACAGCCACAACGTATACATCACTGCTGACAAACAAAAAAACGGCATCAAAGCAAACTTCAAAATCCGTCATAACGTAGAGGACGGTTCTGTTCAGCTTGCTGATCATTATCAGCAAAATACACCGATCGGTGACGGCCCGGTTCTTCTTCCTGATAACCATTATTTATCAACTCAAAGCGTATTATCAAAAGACCCAAATGAAAAGCGTGACCACATGGTGCTGCTTGAATTTGTGACAGCTGCTGGTATCACTCACGGCATGGATGAGCTTTATAAGTAAgGTCTCGGTTTAAGAGCTATGCTGGAAACAGCATAGCAAGTTTAAATAAGGCTAGTCCGTTATCAACTTGAAAAAGTGGCACCGAGTCGGTGCTTTTTTTGAAGCTTGGGCCCGAACAAAAACTCATGGATCC

### Dual CRISPRi library construction

The process of library construction overflow is visualized in Figure S1. Initially, we designed a comprehensive set of 869 sgRNAs for the dual sgRNA library as described in Liu et al 2021^1^, and the complete list of sgRNAs can be found in supplementary Table S1. For each sgRNA, two partially complementary 24-nt oligos were custom-ordered. The cloning of these sgRNAs into the pPEPZ-P3-dualsgRNA vector was performed as previously reported^21^. Specifically, the two oligos were annealed to generate short duplex DNA molecules with 4-nt overhangs on each end, and subsequently phosphorylated. Note that transcription of each sgRNA starts with the +1 of the P3 promoter (adenine)^26^ to ensure equally strong transcription^27^ and prevent undesirable uridines at the 5’ end of the sgRNA^60^. To insert the sgRNAs into the first insertion site, the pPEPZ-P3-dualsgRNA vector underwent BsmBI digestion. The annealed short duplex DNA with sgRNA spacer sequences was ligated to the BsmBI-digested vector using T4 ligase. The resulting ligation product was then introduced into Stbl3 chemical-competent cells (ThermoFisher, Cat. No. C737303) and subsequently selected on LB agar plates containing 100 µg/ml spectinomycin. The colonies displaying green color indicated the successful insertion of sgRNAs at the sgRNA1 insertion site. In our assay, we obtained approximately 22,000 colonies, ensuring a robust 25-fold coverage of the sgRNA library. These colonies were collectively collected and pooled for plasmid isolation. The isolated plasmid served as the vector for the second sgRNA insertion.

The plasmid carrying the sgRNA1 pool was digested with BsaI to enable the cloning of sgRNA2, employing a procedure similar to that used for sgRNA1. The key difference is that the ligation product was introduced into S. pneumoniae instead of *E. coli*, followed by selection on Columbia sheep blood agar plate with 100 µg/ml spectinomycin. This process yielded a total of 9.96 x 10^6^ colonies, providing an extensive 26-fold coverage of the dual sgRNA pool.

#### Sequencing

The barcoded samples were subjected to Illumina NextSeq and NovaSeq6000 sequencing. The NovaSeq600 sequencing was done at the Genomic Technologies Facility in Lausanne, Switzerland. To improve diversity, the samples were mixed with 40% PhiX library. For the NovaSeq6000 run, Read1 was sequenced with 54 dark cycles, followed by 25 standard cycles. The read2 was then sequenced with 115 standard cycles. Both i7 and i5 indexes were sequenced with 8 cycles. All fastq files from sequencing are uploaded to Sequence Read Archive (SRA) on NCBI with accession number PRJNA1105072.

#### Construction of the dual CRISPRi strains

To construct a dual CRISPRi strain, each sgRNA oligo annealing (Figure S1, panel 2) was phosphorylated using T4 PNK, 1mM ATP, and T4 PNK 10x buffer (ThermoFisher, EK0031) and incubated at 37°C for 30 minutes, followed by heat inactivation at 65°C for 20min. Phosphorylated sgRNA oligos were ligated into the vector pVL2118 by successive digestion ligation as described in Figure S1 using Esp3I (isoschizomer of BsmBI, NEB, #R0734S), or BsaI-HF®v2 (NEB, #R3733S), then using T4 DNA ligase and T4 DNA ligase 10x buffer (ThermoFisher, EL0011). Completed plasmid was then isolated from *E. coli* using PureYield™ Plasmid Miniprep (Promega, #TB374), and then transformed into DCI23 (or VL997), and checked by PCR using the 2x Phanta Max Master Mix polymerase (Vazyme®, #P515), using the primers OVL866 and OVL867 (Table S5). The amplified fragment was verified on 1% agarose gel, then purified using a Macherey-Nagel NucleoSpin® Gel and PCR clean-up kit, and checked by Sanger sequencing by Microsynth AG.

### Analysis of the dual CRISPRi sequenced libraries

#### Differential fitness evaluation analysis

To retrieve the read count of each sgRNA combination, we adapted the feature counter python package 2FAST2Q^28^. As the sgRNA in position 1 (SG1) and sgRNA in position 2 (SG2) are individually sequenced in read1 and read2 respectively, we used the read id to merge read1 with read2 and map this to the sgRNA sequences, with a maximum allowance of 3 mismatches. We then counted the number of occurrences of each combination and exported the count table to a csv file. The custom python script is available at https://github.com/veeninglab/Dual-CRISPRi. To compute the fitness cost of each sgRNA combination, the raw counts of each sgRNA combination were analyzed using the R package DESeq2^61^. Since we used two lanes on the NovaSeq6000 flow cell, we aggregated the counts from Lane 1 and Lane 2. For all sgRNA combinations, we tested against an absolute log2FC of 1, with an alpha of 0.05, yielding a log2FC value describing the fitness cost of the combination, between non-induced and induced libraries (CRISPRi OFF vs CRISPRi ON). sgRNA combinations where SG1 and SG2 were the same (e.g. sgRNA137 in position 1, and sgRNA137 in position 2) were used to compute the individual fitness cost of each sgRNA individually. We filtered out sgRNA combinations where induced average read counts was below 4 and expected read counts (calculated using non-induced read count and individual fitness cost of the sgRNA) below 2 reads. This allowed the removal of combinations for which the read counts were already largely depleted in induced conditions, rendering the interaction difficult to interpret. The strength of the genetic interaction was calculated as described by Mani et al. (2008), using an additive model: ε = *Wxy* − *E*(*Wx* + *Wy*), where *Wx* and *Wy* represent the individual fitness of each sgRNA, and *Wxy* the observed fitness of any sgRNA combination^29^. Epsilon values smaller than -1 were grouped as negative interactions, and epsilon values bigger than 1.5 were grouped as positive interactions. For the 19×1498 library, significance of differential sgRNA depletion/enrichment between CRISPRi induction, across each 19 sgRNA using the sgRNA targeting luciferase as a reference fitness evaluation (absolute log2FC > 1, p < 0.05) was tested using DESeq2. The R script used for the analysis is available at https://github.com/veeninglab/Dual-CRISPRi. All plots were created using R (v4.3.3).

### Microplate growth assay

To perform growth assays, the bacterial strains were cultured in fresh C+Y (pH = 6.8) with or without 1 mM IPTG and/or 50 ng/mL aTc at 37°C 5% CO_2_ until reaching OD600 0.2. The cultures were then diluted to OD595 0.01 in fresh C+Y medium (pH = 6.8) with or without 1 mM IPTG and/or 50 ng/mL aTc, and dispensed into a 96-well plate (200 µl per well). Cell growth was monitored by measuring optical density at 595nm every 10 minutes using a microplate reader (TECAN infinite F200 Pro). Each growth assay was performed in triplicate, and the mean value plotted, with the SEM (Standard Error of the Mean) represented by an area around the curve. Each well was normalized by subtracting the minimum value of each well over the period. Area under the curve was calculated from 0h to 10h, in an empirical manner, where OD values are summed over the time. Plots were made using BactEXTRACT^62^.

### Differential AUC fitness calculation

We calculated the empiric area under the curve (AUC) by summing the first 10 hours of growth. We normalized each AUC by the wild type (in which no sgRNA is present) following 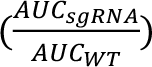. We measured the difference between the growth fitness and the expected growth fitness of the double mutant with a multiplicative model as shown recently, defined by 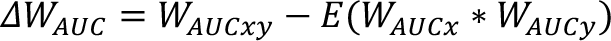^15,29^. We propagated the standard deviation to the expected AUC fitness by 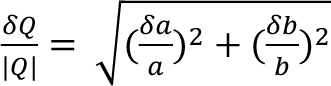, where 0 and 1 are the normalized *W_AUCx_* and *W_AUCx_* values respectively, and where δ*a* and δ*b* are the standard deviations of *W_AUCy_* and *W_AUCy_* respectively.

### Phase contrast Microscopy

Strains stocks were diluted 1:100 into fresh C+Y with necessary inducer (i.e. IPTG, aTc), and incubated at 37°C. When OD_595_ reached 0.15, 1 ml of culture was harvested by centrifugation for 1 min at 9,000*g*. For DAPI staining, cells were resuspended with 1 ml of Phosphate-buffered saline (PBS) and 1 μg/ml of DAPI (Sigma-Aldrich) was added. The Cells were incubated for 5 min at room temperature and then washed two times PBS solution, and resuspended with 200 to 400 μL of PBS. 0.4 μL of the cell suspension was spotted on a thin 1.2% PBS-agarose pad on a microscope slide. Microscopy images were captured using LasX (Leica) DMi8 microscope with a DFC9000 GTC-VSC04862 camera, a HC PL APO 100x/1.40 Oil objective and visualized using sola light engine (Lumencor®). The images were then processed using ImageJ. For DAPI imaging, Leica DMi8 filter was used: Leica 11533333, Ex: 395/25 nm, BS: LP 425 nm, Em: BP 460/50 nm), and image was acquired with an exposure time of 500ms. Images were processed using LasX v.3.4.2.18368 (Leica).

## Acknowledgements

We would like to thank Afonso Bravo and Clement Gallay for help with the analysis and all members of the Veening lab for continued support. We thank the Genomic Technologies Facility in Lausanne for sequencing. Work in the Veening lab is supported by the Swiss National Science Foundation (SNSF) (project grants 310030_192517, 310030_200792 and ‘AntiResist’ 51NF40_180541) and ERC consolidator grant 771534-PneumoCaTChER. Xue Liu was supported by the National Key Research and Development Program of China (2023YFD1800100), the Science and Technology Project of Shenzhen (JCYJ20220818095602006), National Nature Science Foundation of China (82270012), and Shenzhen University 2035 Program for Excellent Research (86901-00000216).

## References

1. de Bakker, V., Liu, X., Bravo, A.M., and Veening, J.W. (2022). CRISPRi-seq for genome-wide fitness quantification in bacteria. Nature Protocols 2021 17:2 17, 252–281. 10.1038/s41596-021-00639-6.

2. Cain, A.K., Barquist, L., Goodman, A.L., Paulsen, I.T., Parkhill, J., and van Opijnen, T. (2020). A decade of advances in transposon-insertion sequencing. Nat Rev Genet 21, 526–540. 10.1038/S41576-020-0244-X.

3. Baryshnikova, A., Costanzo, M., Dixon, S., Vizeacoumar, F.J., Myers, C.L., Andrews, B., and Boone, C. (2010). Synthetic genetic array (SGA) analysis in *Saccharomyces cerevisiae* and *Schizosaccharomyces pombe*. Methods Enzymol 470, 145–179. 10.1016/S0076-6879(10)70007-0.

4. Horlbeck, M.A., Xu, A., Wang, M., Bennett, N.K., Park, C.Y., Bogdanoff, D., Adamson, B., Chow, E.D., Kampmann, M., Peterson, T.R., et al. (2018). Mapping the Genetic Landscape of Human Cells. Cell 174, 953–967.e22. 10.1016/J.CELL.2018.06.010.

5. Costanzo, M., VanderSluis, B., Koch, E.N., Baryshnikova, A., Pons, C., Tan, G., Wang, W., Usaj, M., Hanchard, J., Lee, S.D., et al. (2016). A global genetic interaction network maps a wiring diagram of cellular function. Science (1979) 353. 10.1126/science.aaf1420.

6. Gordon, D.E., Watson, A., Roguev, A., Zheng, S., Jang, G.M., Kane, J., Xu, J., Guo, J.Z., Stevenson, E., Swaney, D.L., et al. (2020). A Quantitative Genetic Interaction Map of HIV Infection. Mol Cell 78, 197. 10.1016/J.MOLCEL.2020.02.004.

7. Babu, M., Arnold, R., Bundalovic-Torma, C., Gagarinova, A., Wong, K.S., Kumar, A., Stewart, G., Samanfar, B., Aoki, H., Wagih, O., et al. (2014). Quantitative Genome-Wide Genetic Interaction Screens Reveal Global Epistatic Relationships of Protein Complexes in *Escherichia coli*. PLoS Genet 10, e1004120. 10.1371/JOURNAL.PGEN.1004120.

8. Babu, M., Díaz-Mejía, J.J., Vlasblom, J., Gagarinova, A., Phanse, S., Graham, C., Yousif, F., Ding, H., Xiong, X., Nazarians-Armavil, A., et al. (2011). Genetic Interaction Maps in *Escherichia coli* Reveal Functional Crosstalk among Cell Envelope Biogenesis Pathways. PLoS Genet 7, e1002377. 10.1371/JOURNAL.PGEN.1002377.

9. Liu, X., Gallay, C., Kjos, M., Domenech, A., Slager, J., van Kessel, S.P., Knoops, K., Sorg, R.A., Zhang, J., and Veening, J. (2017). High-throughput CRISPRi phenotyping identifies new essential genes in *Streptococcus pneumoniae*. Mol Syst Biol 13, 931. 10.15252/msb.20167449.

10. Van Opijnen, T., and Camilli, A. (2012). A fine scale phenotype-genotype virulence map of a bacterial pathogen. Genome Res 22, 2541–2551. 10.1101/gr.137430.112.

11. Rosconi, F., Rudmann, E., Li, J., Surujon, D., Anthony, J., Frank, M., Jones, D.S., Rock, C., Rosch, J.W., Johnston, C.D., et al. (2022). A bacterial pan-genome makes gene essentiality strain-dependent and evolvable. Nature Microbiology 2022 7:10 7, 1580–1592. 10.1038/s41564-022-01208-7.

12. van Opijnen, T., Bodi, K.L., and Camilli, A. (2009). Tn-seq: High-throughput parallel sequencing for fitness and genetic interaction studies in microorganisms. Nat Methods 6, 767–772. 10.1038/nmeth.1377.

13. van Opijnen, T., Lazinski, D.W., and Camilli, A. (2015). Genome-wide fitness and genetic interactions determined by Tn-seq, a high-throughput massively parallel sequencing method for microorganisms. Curr Protoc Microbiol 2015, 1E.3.1-1E.3.24. 10.1002/9780471729259.mc01e03s36.

14. Cain, A.K., Barquist, L., Goodman, A.L., Paulsen, I.T., Parkhill, J., and van Opijnen, T. (2020). A decade of advances in transposon-insertion sequencing. Nature Reviews Genetics 2020 21:9 21, 526–540. 10.1038/s41576-020-0244-x.

15. Jana, B., Liu, X., Dénéréaz, J., Park, H., Leshchiner, D., Liu, B., Gallay, C., Zhu, J., Veening, J.-W., and van Opijnen, T. (2024). CRISPRi-TnSeq maps genome-wide interactions between essential and non-essential genes in bacteria. Nature Microbiology, 10.1038/s41564-024-01759-x.

16. Peters, J.M., Colavin, A., Shi, H., Czarny, T.L., Larson, M.H., Wong, S., Hawkins, J.S., Lu, C.H.S., Koo, B.M., Marta, E., et al. (2016). A Comprehensive, CRISPR-based Functional Analysis of Essential Genes in Bacteria. Cell 165, 1493–1506. 10.1016/J.CELL.2016.05.003.

17. Qi, L.S., Larson, M.H., Gilbert, L.A., Doudna, J.A., Weissman, J.S., Arkin, A.P., and Lim, W.A. (2013). Repurposing CRISPR as an RNA-guided platform for sequence-specific control of gene expression. Cell 152, 1173–1183. 10.1016/j.cell.2013.02.022.

18. Minhas, V., Domenech, A., Synefiaridou, D., Straume, D., Brendel, M., Cebrero, G., Liu, X., Costa, C., Baldry, M., Sirard, J.C., et al. (2023). Competence remodels the pneumococcal cell wall exposing key surface virulence factors that mediate increased host adherence. PLoS Biol 21, e3001990. 10.1371/JOURNAL.PBIO.3001990.

19. Dewachter, L., Dénéréaz, J., Liu, X., Bakker, V. de, Costa, C., Baldry, M., Sirard, J.C., and Veening, J.W. (2022). Amoxicillin-resistant *Streptococcus pneumoniae* can be resensitized by targeting the mevalonate pathway as indicated by sCRilecs-seq. Elife 11. 10.7554/ELIFE.75607.

20. Replogle, J.M., Bonnar, J.L., Pogson, A.N., Liem, C.R., Maier, N.K., Ding, Y., Russell, B.J., Wang, X., Leng, K., Guna, A., et al. (2022). Maximizing CRISPRi efficacy and accessibility with dual-sgRNA libraries and optimal effectors. Elife 11. 10.7554/ELIFE.81856.

21. Han, K., Jeng, E.E., Hess, G.T., Morgens, D.W., Li, A., and Bassik, M.C. (2017). Synergistic drug combinations for cancer identified in a CRISPR screen for pairwise genetic interactions. Nat Biotechnol 35, 463–474. 10.1038/nbt.3834.

22. Ellis, N.A., Kim, B., Tung, J., and Machner, M.P. (2021). A multiplex CRISPR interference tool for virulence gene interrogation in *Legionella pneumophila*. Commun Biol 4. 10.1038/S42003-021-01672-7.

23. Reis, A.C., Halper, S.M., Vezeau, G.E., Cetnar, D.P., Hossain, A., Clauer, P.R., and Salis, H.M. (2019). Simultaneous repression of multiple bacterial genes using nonrepetitive extra-long sgRNA arrays. Nat Biotechnol 37, 1294–1301. 10.1038/S41587-019-0286-9.

24. McCarty, N.S., Graham, A.E., Studená, L., and Ledesma-Amaro, R. (2020). Multiplexed CRISPR technologies for gene editing and transcriptional regulation. Nat Commun 11. 10.1038/S41467-020-15053-X.

25. Liu, X., Kimmey, J.M., Matarazzo, L., de Bakker, V., Van Maele, L., Sirard, J.C., Nizet, V., and Veening, J.W. (2021). Exploration of Bacterial Bottlenecks and *Streptococcus pneumoniae* Pathogenesis by CRISPRi-Seq. Cell Host Microbe 29, 107–120.e6. 10.1016/j.chom.2020.10.001.

26. Sorg, R.A., Kuipers, O.P., and Veening, J.W. (2015). Gene expression platform for synthetic biology in the human pathogen *Streptococcus pneumoniae*. ACS Synth Biol 4, 228–239. 10.1021/SB500229S.

27. Slager, J., Aprianto, R., and Veening, J.-W. (2018). Deep genome annotation of the opportunistic human pathogen *Streptococcus pneumoniae* D39. Nucleic Acids Res, 283663. 10.1093/nar/gky725.

28. Bravo, A.M., Typas, A., and Veening, J.W. (2022). 2FAST2Q: a general-purpose sequence search and counting program for FASTQ files. PeerJ 10, e14041. 10.7717/PEERJ.14041/SUPP-11.

29. Mani, R., St. Onge, R.P., Hartman IV, J.L., Giaever, G., and Roth, F.P. (2008). Defining genetic interaction. Proc Natl Acad Sci U S A 105, 3461–3466. 10.1073/pnas.0712255105.

30. Tsui, H.C.T., Zheng, J.J., Magallon, A.N., Ryan, J.D., Yunck, R., Rued, B.E., Bernhardt, T.G., and Winkler, M.E. (2016). Suppression of a deletion mutation in the gene encoding essential PBP2b reveals a new lytic transglycosylase involved in peripheral peptidoglycan synthesis in *Streptococcus pneumoniae* D39. Mol Microbiol 100, 1039–1065. 10.1111/mmi.13366.

31. Hoskins, J.A., Matsushima, P., Mullen, D.L., Tang, J., Zhao, G., Meier, T.I., Nicas, T.I., and Jaskunas, S.R. (1999). Gene disruption studies of penicillin-binding proteins 1a, 1b, and 2a in *Streptococcus pneumoniae*. J Bacteriol 181, 6552–6555. 10.1128/jb.181.20.6552-6555.1999.

32. Paik, J., Kern, I., Lurz, R., and Hakenbeck, R. (1999). Mutational analysis of the *Streptococcus pneumoniae* bimodular class A penicillin-binding proteins. J Bacteriol 181, 3852–3856. 10.1128/JB.181.12.3852-3856.1999.

33. Fenton, A.K., El Mortaji, L., Lau, D.T.C., Rudner, D.Z., and Bernhardt, T.G. (2017). CozE is a member of the MreCD complex that directs cell elongation in *Streptococcus pneumoniae*. Preprint at NIH Public Access, 10.1038/nmicrobiol.2017.11.

34. Liu, X., Van Maele, L., Matarazzo, L., Soulard, D., Alves Duarte da Silva, V., de Bakker, V., Dénéréaz, J., Bock, F.P., Taschner, M., Ou, J., et al. (2024). A conserved antigen induces respiratory Th17-mediated broad serotype protection against pneumococcal superinfection. Cell Host Microbe 32, 304–314.e8. 10.1016/j.chom.2024.02.002.

35. Butland, G., Babu, M., Díaz-Mejía, J.J., Bohdana, F., Phanse, S., Gold, B., Yang, W., Li, J., Gagarinova, A.G., Pogoutse, O., et al. (2008). eSGA: E. coli synthetic genetic array analysis. Nature Methods 2008 5:9 5, 789–795. 10.1038/nmeth.1239.

36. Lunsford, R.D., and Macrina, F.L. (1986). Molecular cloning and characterization of scrB, the structural gene for the *Streptococcus mutans* phosphoenolpyruvate-dependent sucrose phosphotransferase system sucrose-6-phosphate hydrolase. J Bacteriol 166, 426–434. 10.1128/JB.166.2.426-434.1986.

37. Boulanger, E.F., Sabag-Daigle, A., Baniasad, M., Kokkinias, K., Schwieters, A., Wrighton, K.C., Wysocki, V.H., and Ahmer, B.M.M. (2022). Sugar-Phosphate Toxicities Attenuate *Salmonella* Fitness in the Gut. J Bacteriol 204. 10.1128/JB.00344-22.

38. Minnen, A., Attaiech, L., Thon, M., Gruber, S., and Veening, J.W. (2011). SMC is recruited to oriC by ParB and promotes chromosome segregation in *Streptococcus pneumoniae*. Mol Microbiol 81, 676–688. 10.1111/j.1365-2958.2011.07722.x.

39. Gruber, S. (2018). SMC complexes sweeping through the chromosome: going with the flow and against the tide. Curr Opin Microbiol 42, 96–103. 10.1016/J.MIB.2017.10.004.

40. Stouf, M., Meile, J.C., and Cornet, F. (2013). FtsK actively segregates sister chromosomes in *Escherichia coli*. Proc Natl Acad Sci U S A 110, 11157–11162. 10.1073/PNAS.1304080110/-/DCSUPPLEMENTAL.

41. Stamsås, G.A., Myrbråten, I.S., Straume, D., Salehian, Z., Veening, J.W., Håvarstein, L.S., and Kjos, M. (2018). CozEa and CozEb play overlapping and essential roles in controlling cell division in *Staphylococcus aureus*. Mol Microbiol 109, 615–632. 10.1111/MMI.13999.

42. Stamsås, G.A., Restelli, M., Ducret, A., Freton, C., Garcia, P.S., Håvarstein, L.S., Straume, D., Grangeasse, C., and Kjos, M. (2020). A coze homolog contributes to cell size homeostasis of *streptococcus pneumoniae*. mBio 11, 1–16. 10.1128/mBio.02461-20.

43. Barbuti, M.D., Lambert, E., Myrbråten, I.S., Ducret, A., Stamsås, G.A., Wilhelm, L., Liu, X., Salehian, Z., Veening, J.-W., Straume, D., et al. (2024). The function of CozE proteins is linked to lipoteichoic acid biosynthesis in *Staphylococcus aureus*. mBio 15. 10.1128/MBIO.01157-24.

44. Keller, L.E., Rueff, A.S., Kurushima, J., and Veening, J.W. (2019). Three new integration vectors and fluorescent proteins for use in the opportunistic human pathogen *Streptococcus pneumoniae*. Genes (Basel) 10. 10.3390/genes10050394.

45. Gruber, S., Veening, J.W., Bach, J., Blettinger, M., Bramkamp, M., and Errington, J. (2014). Interlinked Sister Chromosomes Arise in the Absence of Condensin during Fast Replication in *B. subtilis*. Current Biology 24, 293–298. 10.1016/J.CUB.2013.12.049.

46. Mercy, C., Lavergne, J.-P., Slager, J., Ducret, A., Garcia, P.S., Noirot-Gros, M.-F., Dubarry, N., Nourikyan, J., Veening, J.-W., and Grangeasse, C. (2018). RocS drives chromosome segregation and nucleoid occlusion in *Streptococcus pneumoniae*. bioRxiv, 359943. 10.1101/359943.

47. Briggs, N.S., Bruce, K.E., Naskar, S., Winkler, M.E., and Roper, D.I. (2021). The Pneumococcal Divisome: Dynamic Control of *Streptococcus pneumoniae* Cell Division. Preprint at Frontiers Media S.A., 10.3389/fmicb.2021.737396.

48. Le Gouëllec, A., Roux, L., Fadda, D., Massidda, O., Vernet, T., and Zapun, A. (2008). Roles of pneumococcal DivIB in cell division. J Bacteriol 190, 4501–4511. 10.1128/JB.00376-08.

49. Noirclerc-Savoye, M., Le Gouëllec, A., Morlot, C., Dideberg, O., Vernet, T., and Zapun, A. (2005). In vitro reconstitution of a trimeric complex of DivIB, DivIC and FtsL, and their transient co-localization at the division site in *Streptococcus pneumoniae*. Mol Microbiol 55, 413–424. 10.1111/J.1365-2958.2004.04408.X.

50. Marmont, L.S., and Bernhardt, T.G. (2020). A conserved subcomplex within the bacterial cytokinetic ring activates cell wall synthesis by the FtsW-FtsI synthase. Proc Natl Acad Sci U S A 117, 23879–23885. 10.1073/PNAS.2004598117/-/DCSUPPLEMENTAL.

51. Barendt, S.M., Land, A.D., Sham, L.T., Ng, W.L., Tsui, H.C.T., Arnold, R.J., and Winkler, M.E. (2009). Influences of capsule on cell shape and chain formation of wild-type and pcsB mutants of serotype 2 *Streptococcus pneumoniae*. J Bacteriol 191, 3024–3040. 10.1128/JB.01505-08.

52. Weiser, J.N., Ferreira, D.M., and Paton, J.C. (2018). *Streptococcus pneumoniae*: transmission, colonization and invasion. Nat Rev Microbiol 16, 355. 10.1038/S41579-018-0001-8.

53. Shirley, J.D., Nauta, K.M., Gillingham, J.R., Diwakar, S., and Carlson, E.E. (2024). kinact/KI Value Determination for Penicillin-Binding Proteins in Live Cells. bioRxiv, 2024.05.05.592586. 10.1101/2024.05.05.592586.

54. van Opijnen, T., Bodi, K.L., and Camilli, A. (2009). Tn-seq: High-throughput parallel sequencing for fitness and genetic interaction studies in microorganisms. Nat Methods 6, 767–772. 10.1038/nmeth.1377.

55. Pang, T., Wang, X., Lim, H.C., Bernhardt, T.G., and Rudner, D.Z. (2017). The nucleoid occlusion factor Noc controls DNA replication initiation in *Staphylococcus aureus*. PLoS Genet 13, 1–26. 10.1371/journal.pgen.1006908.

56. Pinho, M.G., Kjos, M., and Veening, J.W. (2013). How to get (a)round: Mechanisms controlling growth and division of coccoid bacteria. Nat Rev Microbiol 11, 601–614. 10.1038/nrmicro3088.

57. Wu, L.J., and Errington, J. (2004). Coordination of cell division and chromosome segregation by a nucleoid occlusion protein in *Bacillus subtilis*. Cell 117, 915–925. 10.1016/j.cell.2004.06.002.

58. Cacace, E., Kim, V., Knopp, M., Tietgen, M., Brauer-Nikonow, A., Inecik, K., Mateus, A., Milanese, A., Mårli, M.T., Mitosch, K., et al. (2022). High-throughput profiling of drug interactions in Gram-positive bacteria. bioRxiv, 2022.12.23.521747. 10.1101/2022.12.23.521747.

59. Gallay, C., Sanselicio, S., Anderson, M.E., Soh, Y.M., Liu, X., Stamsås, G.A., Pelliciari, S., van Raaphorst, R., Dénéréaz, J., Kjos, M., et al. (2021). CcrZ is a pneumococcal spatiotemporal cell cycle regulator that interacts with FtsZ and controls DNA replication by modulating the activity of DnaA. Nat Microbiol 6, 1175– 1187. 10.1038/s41564-021-00949-1.

60. van Gestel, J., Hawkins, J.S., Todor, H., and Gross, C.A. (2021). Computational pipeline for designing guide RNAs for mismatch-CRISPRi. STAR Protoc 2. 10.1016/J.XPRO.2021.100521.

61. Love, M.I., Huber, W., and Anders, S. (2014). Moderated estimation of fold change and dispersion for RNA-seq data with DESeq2. Genome Biol 15, 550. 10.1186/s13059-014-0550-8.

62. Dénéréaz, J., and Veening, J.-W. (2024). BactEXTRACT: an R Shiny app to quickly extract, plot and analyse bacterial growth and gene expression data. Access Microbiol 6, 000742.v3. 10.1099/ACMI.0.000742.V3.

## References

1. Mani, R., St. Onge, R.P., Hartman IV, J.L., Giaever, G., and Roth, F.P. (2008). Defining genetic interaction. Proc Natl Acad Sci U S A 105, 3461–3466. 10.1073/pnas.0712255105.

2. Liu, X., Kimmey, J.M., Matarazzo, L., de Bakker, V., Van Maele, L., Sirard, J.C., Nizet, V., and Veening, J.W. (2021). Exploration of Bacterial Bottlenecks and Streptococcus pneumoniae Pathogenesis by CRISPRi-Seq. Cell Host Microbe 29, 107–120.e6. 10.1016/j.chom.2020.10.001.

3. Keller, L.E., Rueff, A.S., Kurushima, J., and Veening, J.W. (2019). Three new integration vectors and fluorescent proteins for use in the opportunistic human pathogen Streptococcus pneumoniae. Genes (Basel) 10. 10.3390/genes10050394.

4. Liu, X., Gallay, C., Kjos, M., Domenech, A., Slager, J., van Kessel, S.P., Knoops, K., Sorg, R.A., Zhang, J., and Veening, J. (2017). High-throughput CRISPRi phenotyping identifies new essential genes in *Streptococcus pneumoniae*. Mol Syst Biol 13, 931. 10.15252/msb.20167449.

