## Supplemental figures for "Dual CRISPRi-Seq for genome-wide genetic interaction studies identifies key genes involved in the pneumococcal cell cycle"

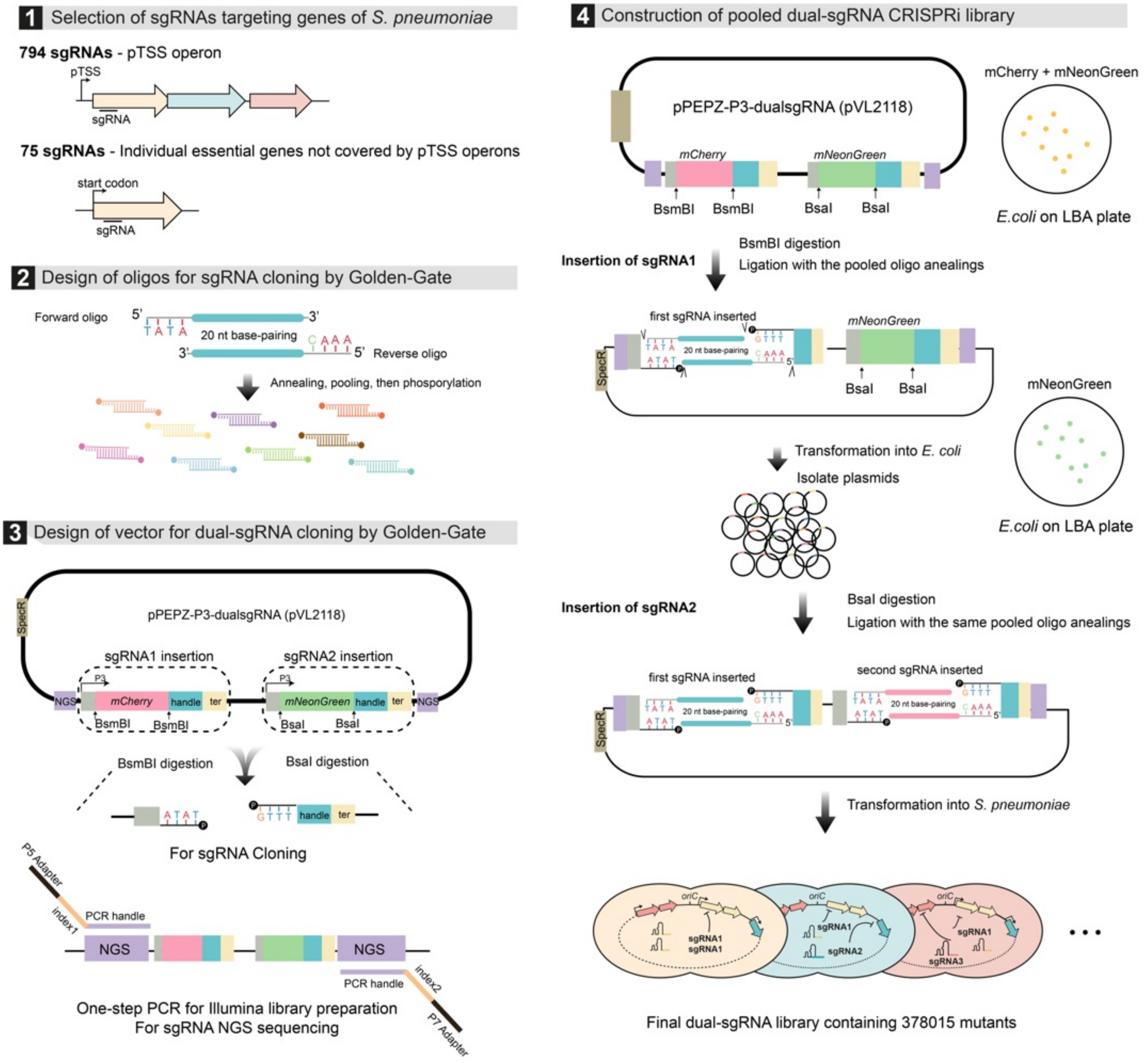


**Figure S1. Dual CRISPRi library construction**. **(1)** A total of 869 sgRNAs were designed for the dual sgRNA library[1]. **(2)** Two oligos were designed for each sgRNA and annealed before Golden Gate Assembly into the vector. **(3)** The pPEPZ-P3-dualsgRNA vector was specifically engineered for dual sgRNA cloning. This vector features two sgRNA insertion sites flanked by BsmBI or BsaI recognition sites, respectively. The coding sequences of mCherry and mNeonGreen were inserted at the cloning sites for sgRNA1 and sgRNA2, respectively. Additionally, DNA sequences for Illumina sequencing, denoted as "NGS," were added to both ends of the dual sgRNA sequence. This design allows for a one-step PCR during Illumina amplicon library construction. **(4)** *E. coli* transformed with the pPEPZ-P3-dualsgRNA vector exhibit orange colonies, a result of simultaneous expression of the red fluorescent protein mCherry and the green fluorescent protein GFP. The initial sgRNA was integrated into the vector through BsmBI digestion and ligation. Upon transformation into *E. coli*, the colonies that exhibited green coloration confirmed successful integration, indicating the replacement of mCherry with sgRNA1. The second sgRNA was incorporated into the vector via BsaI digestion and ligation, after which the construct was transformed into *S. pneumoniae* to construct the final dual CRISPRi library.

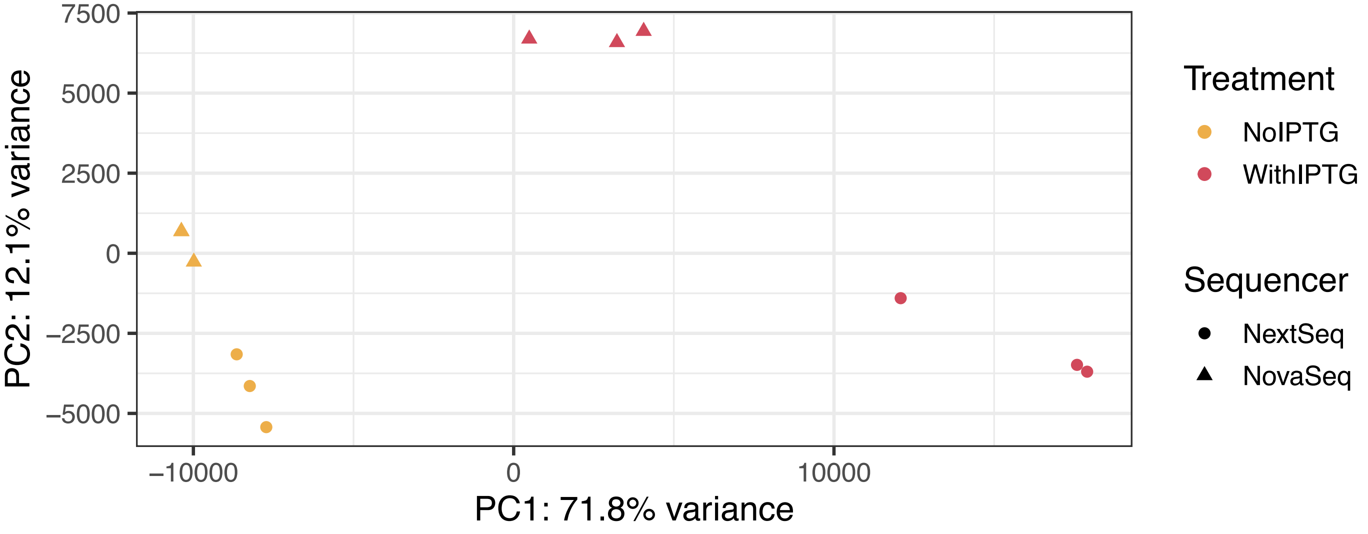


**Figure S2. PCA of all samples.** Principal component analysis (PCA) shows the disparity between the samples and the treatment. Orange points represent samples where CRISPRi was activated by IPTG, and red points show samples where CRISPRi was not activated. The treatment explains most of the variance in sample strain composition.

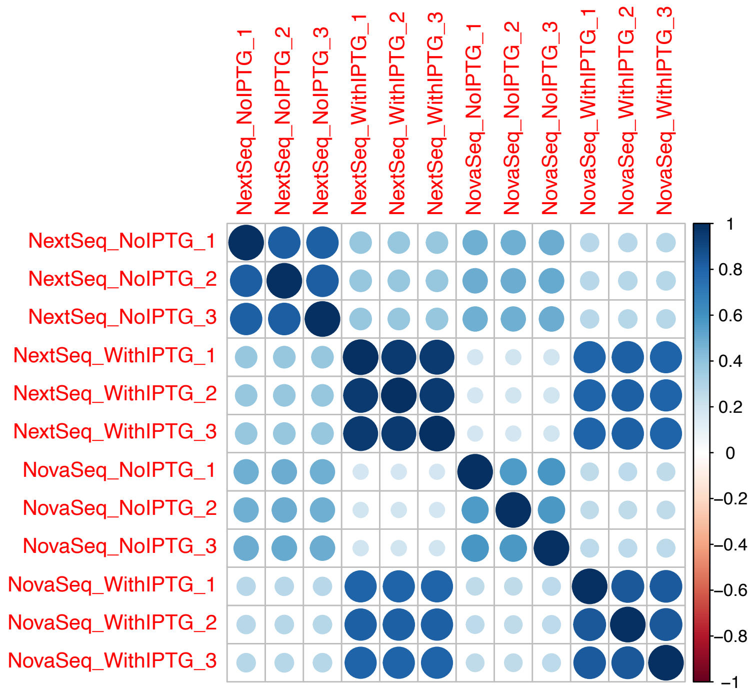


**Figure S3. Pairwise correlations between all samples.** Pairwise Pearson correlations were computed between all replicates. Dark blue shows a positive correlation, and dark red shows a negative correlation. High correlations between equal treatments were observed.

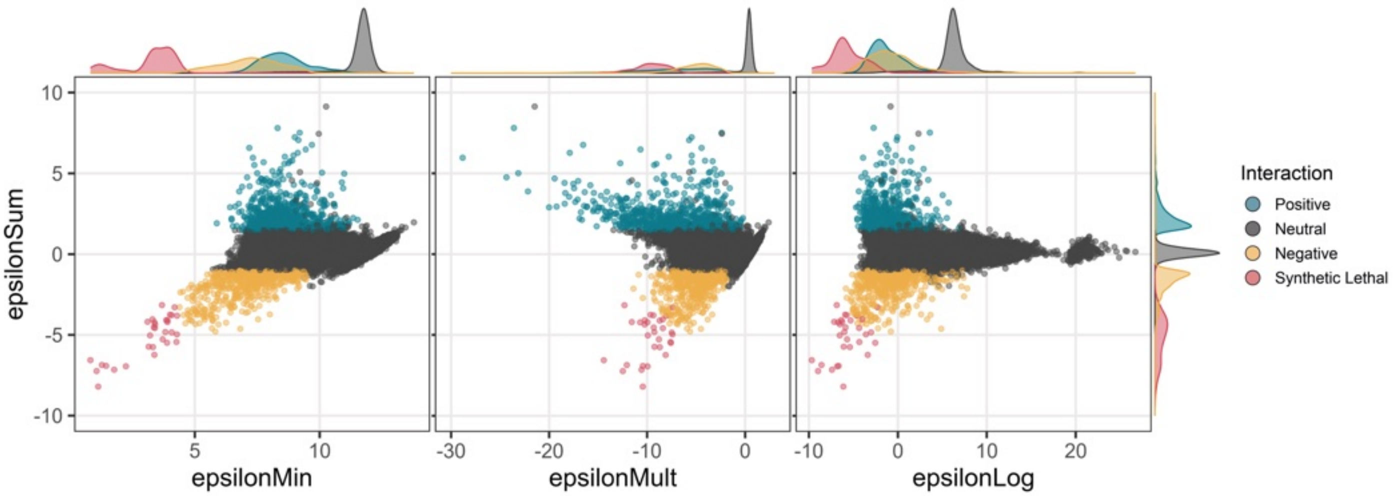


**Figure S4. Epsilon score models.** Distribution comparison between sum model $\varepsilon = Wxy - E(Wx + Wy)$, with either minimum model $\varepsilon = Wxy - \min(Wx, Wy)$, multiplicative model $\varepsilon = Wxy - E(Wx * Wy)$, or logarithmic model $\varepsilon = Wxy - log2((2^{Wx} - 1) * (2^{Wy} - 1))$ as described before[2]. Each sgRNA combination is shown as a point. The color of the point represents the identified genetic interaction using the sum model. The density of each population is shown in the margins of each plot.

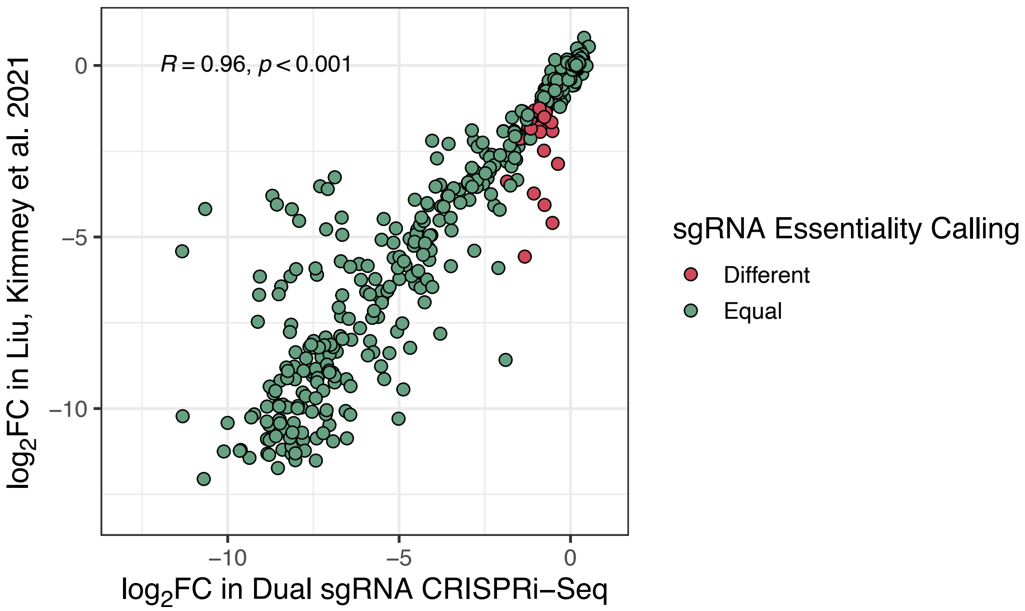


**Figure S5. Correlation with fitness scores of Liu et al.** We compared the log2FC values obtained in the dual sgRNA library for the individual 869 sgRNAs with the log2FC obtained for these same 869 sgRNAs of Liu, Kimmey et al.[1], in the same growth conditions. The Pearson correlation was calculated and shown. The green points represent when the sgRNA essentiality calling is equal between the two studies. sgRNA essentiality is defined a significantly different normalized abundance upon activation of CRISPRi ($log_2FC < -1$, $padj < 0.05$).

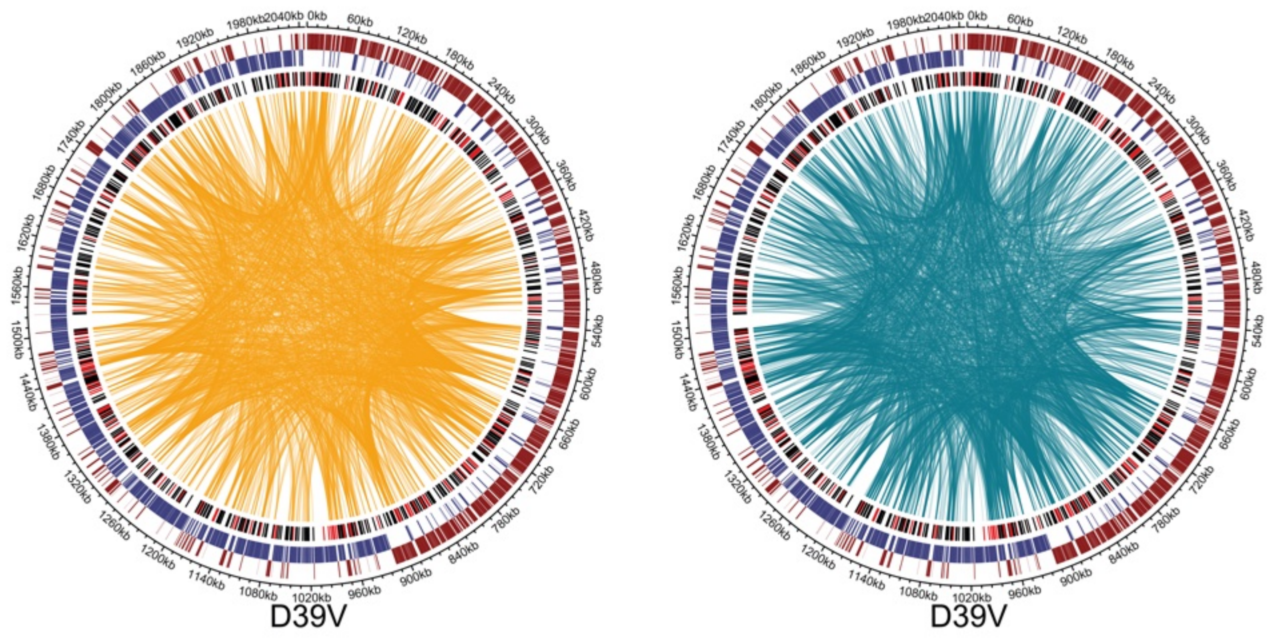


**Figure S6. CIRCOS plots representing all genetic interactions identified.** From outer ring to inner ring: D39V genome position, genes (dark red) on the leading positive strand, genes (dark blue) located on the negative strand, position of sgRNA targets (red if the targeted gene(s) are essential, black if the targeted gene(s) are not essential). Each genetic interaction is represented by a link between 2 sgRNAs. Orange for negative genetic interaction (left CIRCOS plot) and blue for positive genetic interaction (right CIRCOS plot).

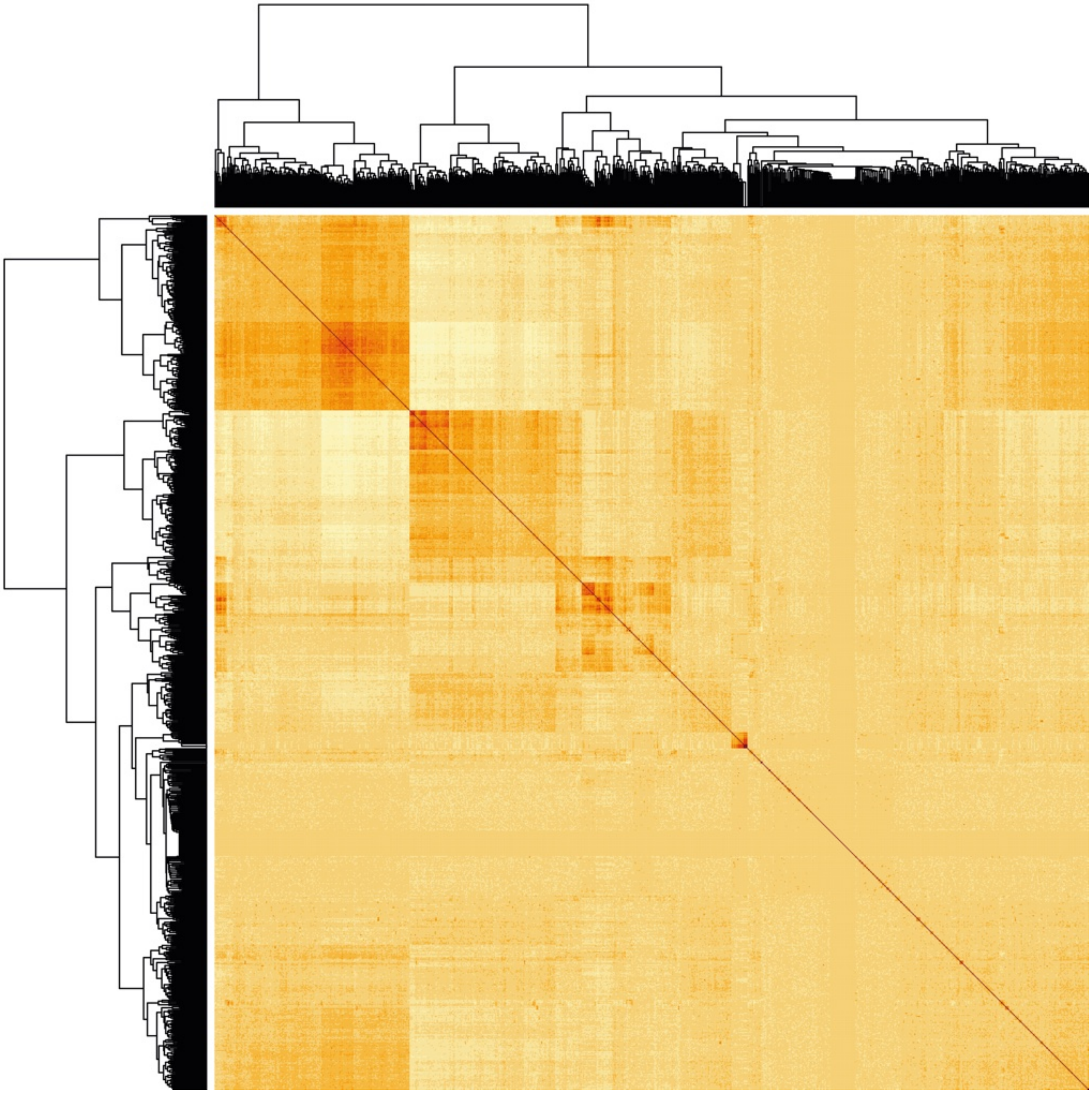

**Figure S7. Heatmap.** Clustered correlation heatmap of all combinations of the 869x869 dual library. The white (0) to red (1) gradient shows the Pearson correlation of Epsilon values between every sgRNA combinations.

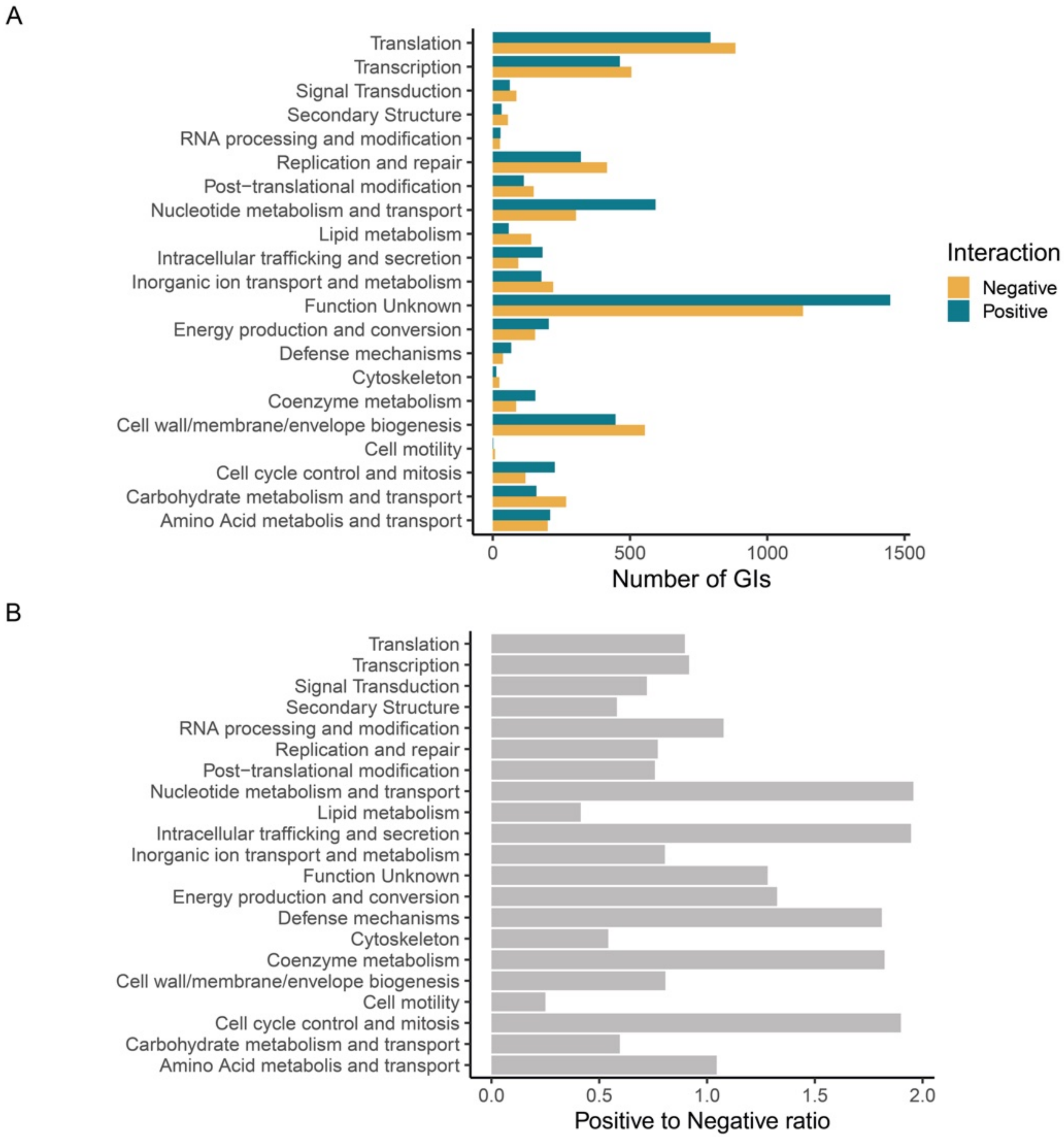


**Figure S8. Clusters of orthologs groups (COG) analysis. A.** Number of GIs per COG category**. B**. Ratios of the number of positive to negative genetic interactions. For each COG, the ratio was calculated of the number of positive interactions over the number of negative interactions.

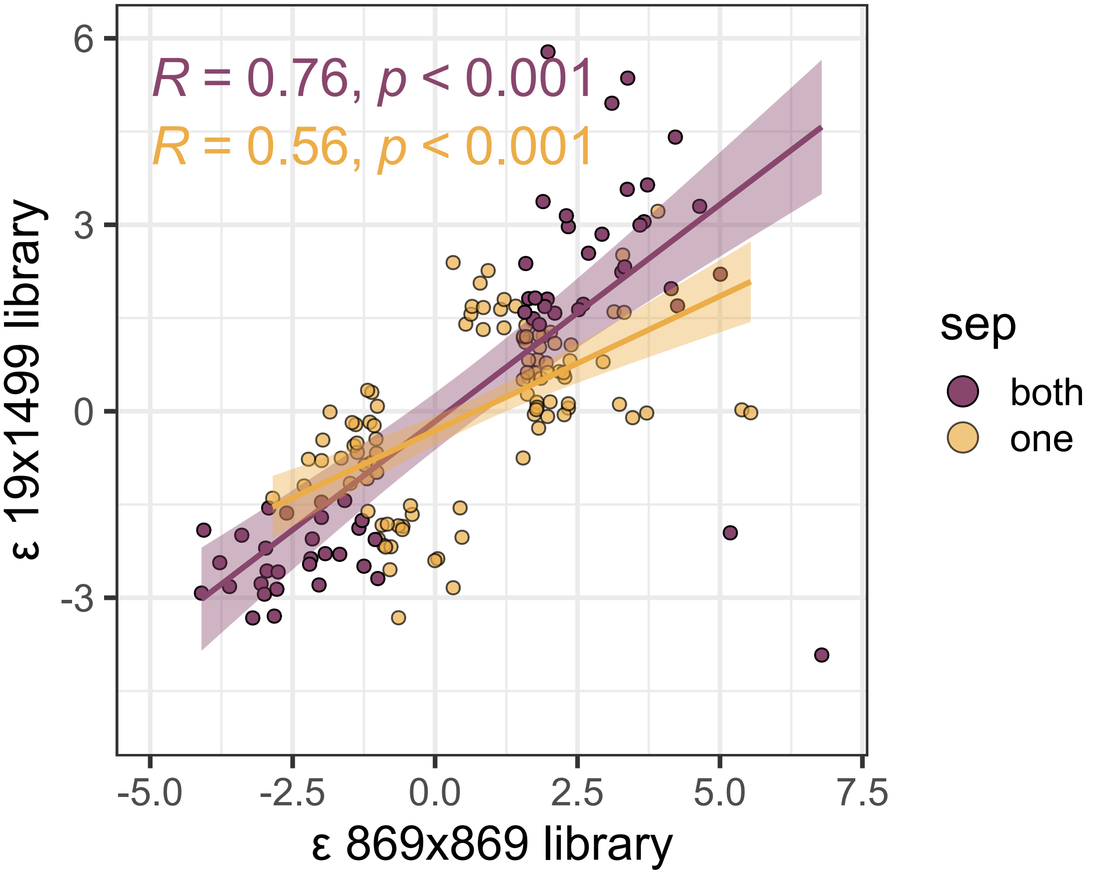


**Figure S9. Correlation of the log2FC fitness scores of the mini and full CRISPRi libraries.** Only the sgRNA combinations covered by both libraries and with a significant interaction in at least one library are shown. Purple points indicate that the genetic interaction was determined to be significant in both libraries, while yellow points indicate that the significance of the genetic interaction was only observed in one library. Pearson correlations and linear regression with confidence intervals are shown.

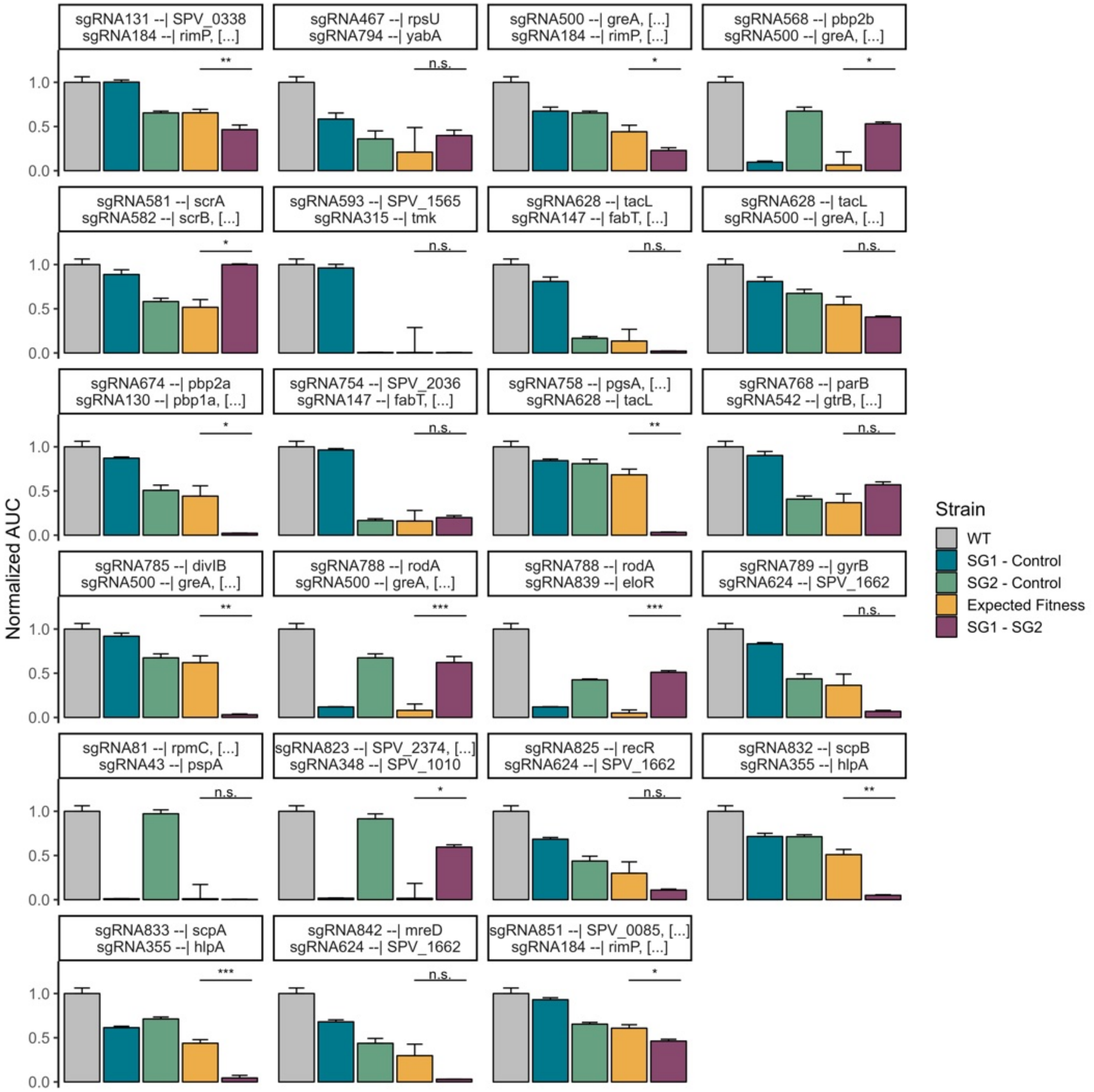


**Figure S10. Area under the curve (AUC) bar plots of dual CRISPRi strains.** AUC was calculated for the first 10 hours of growth. Each facet corresponds to a specific sgRNA combination, with sgRNA targets shown in the facet title (left corresponds to gene(s) targeted by SG1, right to gene(s) targeted by SG2). Either no sgRNA present (grey, WT), only SG1 present (blue), only SG2 present (green), or both sgRNA present (purple). Presence of “[…]” in the facets title means the existence of additional genes in the same operon which can also be targeted by the same sgRNA (See Table S1). Every AUC was normalized by the WT, and the expected fitness was calculated based on a multiplicative model (See methods). T-tests were performed to compare the expected fitness (yellow) with the observed fitness (purple). *** means $p < 0.001$, ** means $p < 0.01$, * means $p < 0.05$, n.s. means non-significant. Standard deviation of the triplicate is shown as error bar.

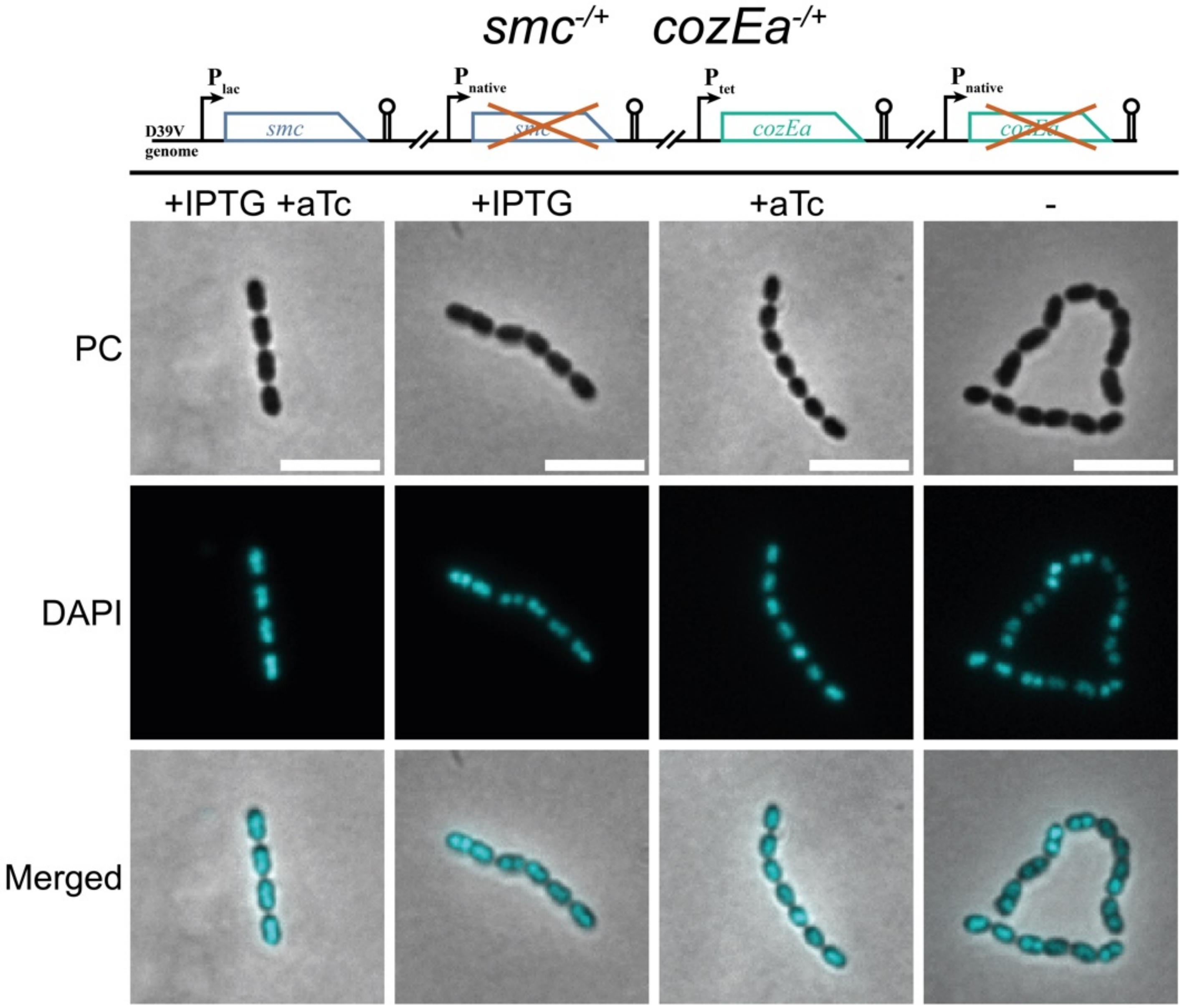


**Figure S11. Morphology of SMC and CozEa depletion mutants.** The s*mc* and *cozEa* double depletion was made by ectopic expression via IPTG-inducible promoter Plac for *smc*, or aTc-inducible promoter for *cozEa* (VL4072, Table S4). 1mM IPTG was added for expression of *smc*, and 50ng/ml aTc was added for induction of *cozEa*. Phase contrast, DAPI staining, and merged microscopy images are shown. Scale bars in white are 5 μm.

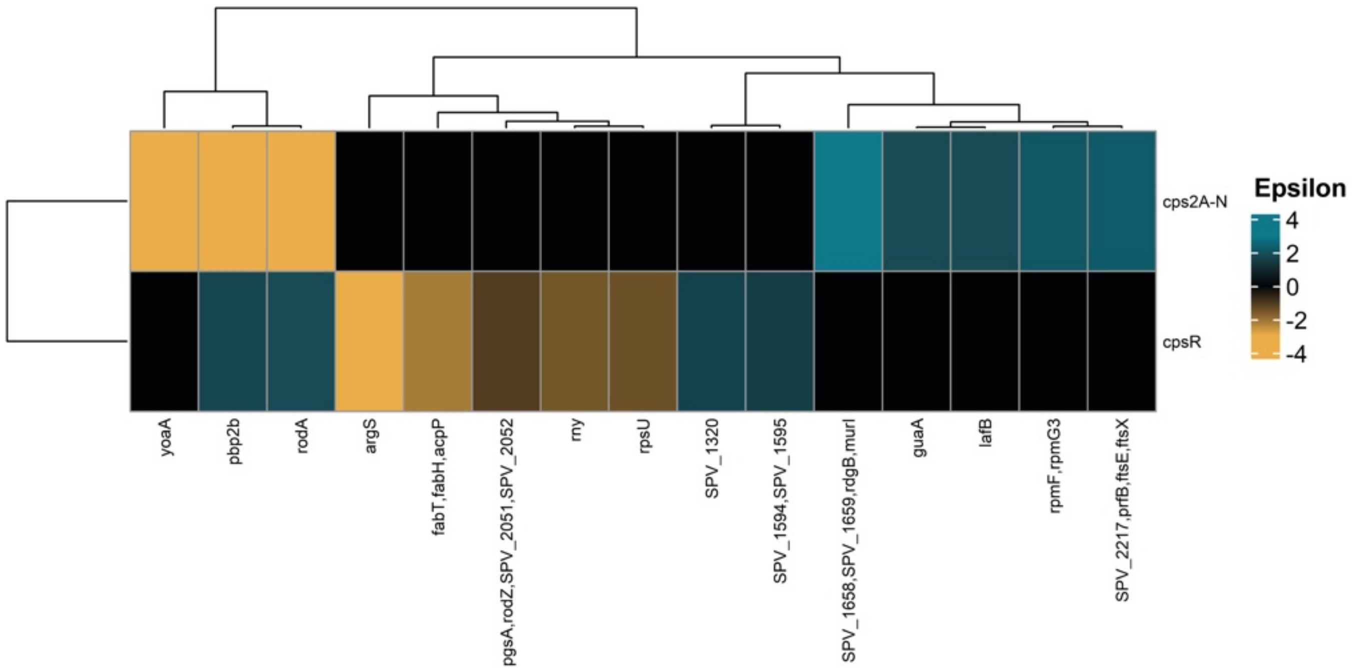


**Figure S12. Clustered heatmap of significant genetic interactions involving either *cps2A-N* or *cpsR* operons.** The blue-to-yellow gradient indicates the strength of genetic interaction, ranging from positive to negative.

### Supplementary references


1. Liu, X., Kimmey, J.M., Matarazzo, L., de Bakker, V., Van Maele, L., Sirard, J.C., Nizet, V., and Veening, J.W. (2021). Exploration of Bacterial Bottlenecks and *Streptococcus pneumoniae* Pathogenesis by CRISPRi-Seq. Cell Host Microbe *29*, 107-120.e6. https://doi.org/10.1016/j.chom.2020.10.001.

2. Mani, R., St. Onge, R.P., Hartman IV, J.L., Giaever, G., and Roth, F.P. (2008). Defining genetic interaction. Proc Natl Acad Sci U S A *105*, 3461–3466. https://doi.org/10.1073/pnas.0712255105.
